# Ninein domains required for its localization, association with partners dynein and ensconsin, and microtubule organization

**DOI:** 10.1101/2023.06.22.546109

**Authors:** Marisa M. L. Tillery, Chunfeng Zheng, Yiming Zheng, Timothy L. Megraw

## Abstract

Ninein (Nin) is a microtubule (MT) anchor at the subdistal appendages of mother centrioles and the pericentriolar material (PCM) of centrosomes that also functions to organize microtubules at non-centrosomal microtubule-organizing centers (ncMTOCs). In humans, the *NIN* gene is mutated in Seckel syndrome, an inherited developmental disorder. Here we dissect the protein domains involved in Nin’s localization and interactions with dynein and ensconsin (ens/MAP7) and show that the association with ens cooperatively regulates microtubule assembly in *Drosophila* fat body cells. We define domains of Nin responsible for its localization to the ncMTOC on the fat body cell nuclear surface, localization within the nucleus, and association with Dynein light intermediate chain (Dlic) and ens, respectively. We show that Nin’s association with ens synergistically regulates MT assembly. Together, these findings reveal novel features of Nin function and its regulation of a ncMTOC.

## Introduction

Ninein (Nin) is a microtubule (MT)-anchoring protein that localizes to subdistal appendages on the mother centriole, within the pericentriolar material (PCM) at the centrosome, and at subcellular sites where non-centrosomal microtubule-organizing centers (ncMTOCs) are organized (Bouckson-Castaing *et al*., 1996; Mogensen *et al*., 2000; Dammermann and Merdes, 2002; Ou *et al*., 2002; Delgehyr *et al*., 2005; Dyachuk *et al*., 2016; Kowanda *et al*., 2016; Zheng *et al*., 2016; Muroyama and Lechler, 2017; Sanchez and Feldman, 2017; Wu and Akhmanova, 2017; Paz and Luders, 2018; Tillery *et al*., 2018; Vineethakumari and Luders, 2022). Mutations in *NIN*, also known as *SCKL7*, are linked to Seckel syndrome, a type of congenital microcephalic primordial dwarfism disorder (Dauber *et al*., 2012). Underscoring its role in growth and development, Nin is essential in neural progenitor cells where it is needed for the asymmetric segregation of mother and daughter centrosomes (Wang *et al*., 2009) and for cell cycle-dependent nuclear movement and MT aster formation at centrosomes (Shinohara *et al*., 2013). During epidermal progenitor cell division, Nin is required for mitotic spindle orientation. A *Nin* null mutation is semi-lethal in mice, and survivors show disruption of desmosomes and lamellar body secretion in keratinocytes resulting in a thin skin phenotype (Lecland *et al*., 2019).

Ninein-like protein (Nlp), a Nin paralog, is also a centrosomal protein (Casenghi *et al*., 2003) with indirect links to ciliopathies (van Wijk *et al*., 2009). Like Nin, Nlp associates with dynein (Redwine *et al*., 2017) and *γ*-tubulin (Casenghi *et al*., 2003). Nlp was shown to be an essential component of the antiviral innate immune response. *NINL* human knockout cells showed enhanced viral replication, making them more susceptible to infection (Stevens *et al*., 2022). In contrast, *Drosophila* has just one *Nin* ortholog, and *Nin* null mutants are viable and fertile (Kowanda *et al*., 2016; Zheng *et al*., 2016). Despite these findings establishing the importance of Nin and Nlp in health and development, little is understood about the molecular functions of Nin and how it organizes an MTOC.

In addition to its localization at centrosomes, Nin is also a component of ncMTOCs where it also functions as a MT anchor (Mogensen *et al*., 2000; Casenghi *et al*., 2003; Delgehyr *et al*., 2005; Wang *et al*., 2015; Zheng *et al*., 2016; Sanchez and Feldman, 2017; Tillery *et al*., 2018). Examples of Nin’s involvement at ncMTOCs include localizing apically in mammalian cochlear cells (Mogensen *et al*., 2000), at the cell cortex in the murine epidermis (Lechler and Fuchs, 2007), and perinuclearly in mammalian and *Drosophila* myotubes (Bugnard *et al*., 2005; Rosen *et al*., 2019) and *Drosophila* larval fat body cells (Zheng *et al*., 2020). In differentiating keratinocytes, Nin is necessary for the cortical organization of MTs and the relocalization of MT-organizing proteins to the cell cortex (Lecland *et al*., 2019). In epithelial cells, the development of an apical-basal polar array of MTs involves a switch from centrosomal MTs that requires Nin and its trafficking by CLIP-170 (Goldspink *et al*., 2017). In the developing vasculature, Nin is required to control tubular morphogenesis of angiogenic endothelial cells (Matsumoto *et al*., 2008). In the mouse brain, alternative splicing of the *Nin* transcript results in expression of a non-centrosomal isoform, implicating a non-centrosomal role in neurons as well (Zhang *et al*., 2016).

Functional requirements for Nin at ncMTOCs are emerging. When centrioles are experimentally eliminated from interphase human cell culture, a single acentriolar MTOC forms from the assembly of PCM proteins CDK5RAP2, Pericentrin, Nin, and *γ*-tubulin. Nin is required for assembly of this ncMTOC, acting late in the assembly process to promote a coalescence of smaller PCM clusters into a compact MTOC and for the formation of the radial MT network (Chen *et al*., 2022).

In *Caenorhabditis elegans* larval epidermis, the *Nin* ortholog *NOCA-1* functions in parallel with the MT stabilizer and Patronin/CAMSAP ortholog *PTRN-1* to organize a ncMTOC critical for growth and morphogenesis. NOCA-1 works with γ-tubulin while PTRN-1 does not (Wang *et al*., 2015). And in *C. elegans* neurons, dendritic ncMTOCs on RAB-11-positive vesicles require NOCA-1 and PTRN-1 (He *et al*., 2022).

In *Drosophila* embryonic muscle, Nin is not essential for nuclear positioning, but its loss sensitizes cells to heterozygous loss of *ens* to affect nuclear positioning. Additionally, Nin overexpression impairs nuclear positioning, but co-overexpression of ens suppresses this (Rosen *et al*., 2019). In *Drosophila* larval fat body cells, a perinuclear MTOC requires the parallel activities of Nin and Patronin (Zheng *et al*., 2020). The Nesprin Muscle-specific protein 300 kDa (Msp300) organizes the fat body perinuclear MTOC by recruiting Patronin and the MT polymerase mini spindles (msps) (Zheng *et al*., 2020). This ncMTOC controls MT assembly, nuclear positioning, and retrograde endocytic trafficking (Zheng *et al*., 2020). The role of Nin at the fat body MTOC, however, is unclear. Although progress has been made in understanding Nin’s localization to various MTOCs and its roles in cell function, development, and disease, we lack a clear understanding of how Nin functions in MT organization and MTOC control.

Human NIN binds to dynein motor (Redwine *et al*., 2017; Celestino *et al*., 2019; Lee *et al*., 2020); *Drosophila* Nin binds to MTs (Kowanda *et al*., 2016) and associates with ensconsin (ens/MAP7) (Rosen *et al*., 2019); and human, *Drosophila*, and *C. elegans* Nin orthologs all associate with *γ*-tubulin (Casenghi *et al*., 2003; Delgehyr *et al*., 2005; Lin *et al*., 2006; Wang *et al*., 2015; Zheng *et al*., 2016). It is likely that these interactions are conserved across species that express Nin orthologs. In *Drosophila* embryonic muscle, Nin interacts and colocalizes with ens to control myonuclear positioning (Rosen *et al*., 2019). Nin and Nlp associate with the dynein-dynactin complex and are activating dynein adapters (Redwine *et al*., 2017; Reck-Peterson *et al*., 2018; Celestino *et al*., 2019) that bind directly to DLIC (Celestino *et al*., 2019; Lee *et al*., 2020). Nin transport to the centrosome is dynein-dependent as inhibiting dynein through excess p50 (Dynamitin) or p150^glued^ CC1 causes a reduction of Nin at the centrosome (Dammermann and Merdes, 2002; Casenghi *et al*., 2005), and Nin may be trafficked along MTs via the dynein complex (Moss *et al*., 2007). Furthermore, loss of NINL leads to a reduction in dynein-dependent transport of intracellular cargoes (Stevens et al., 2022). Currently, the functional relationships between Nin and its partners are poorly understood.

To understand how Nin functions in MT assembly, we investigated the respective contributions its protein domains make to its localization at the nuclear surface and its relationships with ens and dynein. These genetic and cell biological findings indicate that multiple domains contribute to Nin localization to the nuclear surface while a central domain is responsible for localizing Nin inside the nucleus. Furthermore, we confirm that *Drosophila* Nin binds to Dlic through the N-terminus of Nin. And finally, we map the ens-binding domain to Nin and show that Nin cooperates with ens to organize MTs.

## Results

### Localization of Nin to the MTOC involves N- and C-terminal domains

We conducted a structure-function analysis of Nin (Figure 1) in the *Drosophila* larval fat body, a tissue analogous to human liver or adipocytes that features large, monolayered cells with a perinuclear ncMTOC (Zheng *et al*., 2020). We generated a series of transgenic constructs that express regions of Nin fused with C-terminal TagRFP and Myc tags (Figure 1B). We based the design of these constructs on previously-annotated domains of Nin that include the N-terminal *γ*-tubulin-binding (Casenghi *et al*., 2003; Delgehyr *et al*., 2005; Zheng *et al*., 2016), MT-binding (Kowanda *et al*., 2016), and dynein-binding region of Nin (Redwine *et al*., 2017; Celestino *et al*., 2019) and a central region of Nin that binds ens (Rosen *et al*., 2019). Taking advantage of the *Drosophila* UAS-GAL4 binary expression system, we individually expressed these transgenes (Figure 1) using SPARC-GAL4, a fat body driver, to evaluate their subcellular localizations in fat body cells (Figure 2, Supplementary Figures 1A-B and 2). A co-IP of full-length Nin (Nin^1-1091^-TagRFP-Myc) with Nin-GFP indicated that Nin can multimerize (Supplementary Figure 1C), so we compared Nin localization in a *Nin*^*1*^ null mutant background (Figure 2) to a wild-type background (Supplementary Figure 1A-B). Localization patterns, however, did not differ whether endogenous Nin was absent (Figure 2) or present (Supplementary Figure 1A-B).

**Figure 1.**
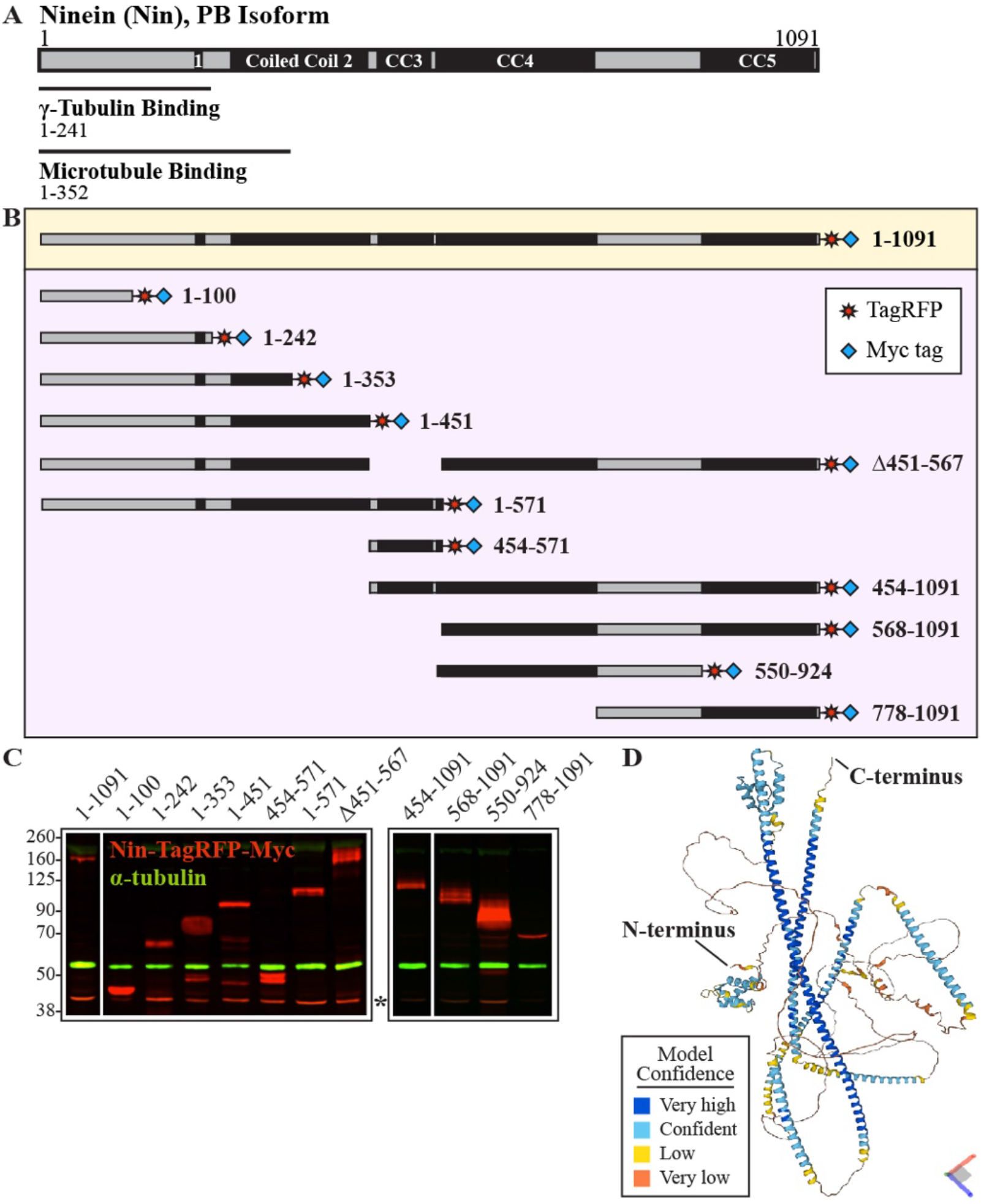
Ninein transgenic constructs for *in vivo* structure-function analysis. (A) The N-terminal half of the 1091-amino acid *Drosophila* Ninein (Nin) protein (isoform B) showing regions that bind *γ*-tubulin (aa 1-241) and microtubules (aa 1-352). Solid black boxes represent coiled coil (CC) regions predicted by COILS software. (B) A full-length, transgenic construct (yellow box) in addition to eleven other constructs that divide Nin into various domains (purple box) for structure-function analysis. Constructs were tagged with TagRFP (red star) and Myc tags (blue diamond) at the C-terminus. Numbers reflect amino acid positions of Nin-PB isoform. (C) Western blot of whole larval lysates probed with an antibody against Myc (red). Nin transgenes driven with SPARC-GAL4 express proteins of molecular weights consistent with their predicted sizes: 1-1091 = 156 kDa; 1-100 = 43.6 kDa; 1-242 = 59.2 kDa; 1-353 = 71.7 kDa; 1-451 = 83.4 kDa; 454-571 = 46.2 kDa; 1-571 = 97.1 kDa; Δ451-567 = 142.8 kDa; 454-1091 = 105.2 kDa; 568-1091 = 92.3 kDa; 550-924 = 74.1 kDa; and 778-1091 = 67.9 kDa. α-Tubulin (green, 50 kDa) served as a loading control. Asterisk denotes a non-specific band. Black frames indicate lanes that were run on a single gel. (D) Predicted structure of Nin (isoform B) from AlphaFold (Jumper *et al*., 2021; Varadi *et al*., 2022).

**Figure 2.**
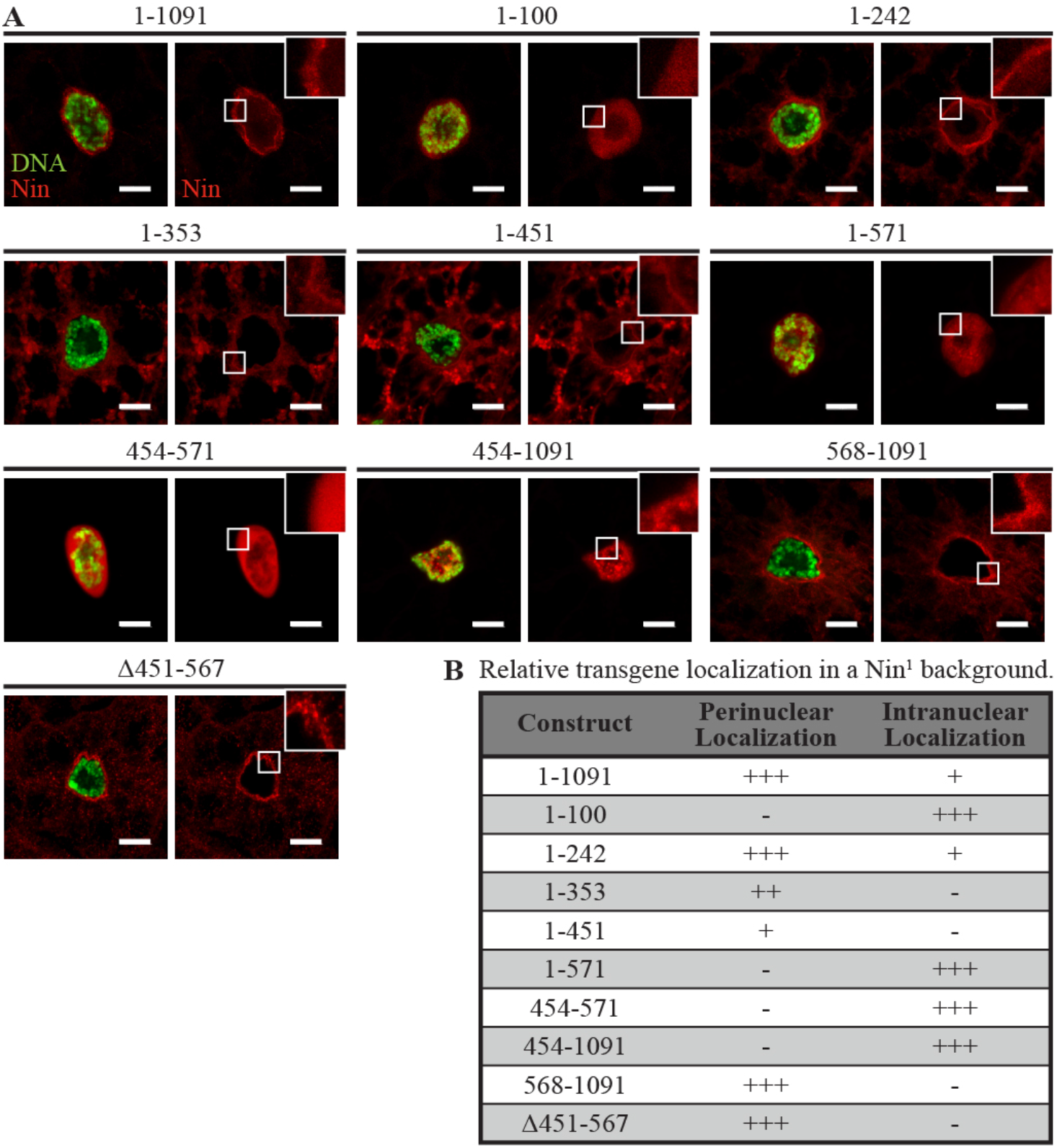
Multiple domains localize Ninein to the nuclear surface or inside the nucleus. The localization patterns in a *Nin* null mutant are shown. IF staining of DNA (DAPI, green) in larval fat body cells expressing Nin transgenic proteins (TagRFP fluorescence, red). Scale bar = 10 μm in this and all subsequent figures. Insets show a magnified view of the nuclear surface. (A) Localization patterns of TagRFP-Myc-labelled Nin transgenes expressed in *Nin*^*1*^ null mutant fat body cells. Multiple domains of Nin, detected with TagRFP fluorescence, localize to the MTOC. Constructs containing amino acids 454-567 localize intranuclearly. (B) Summary of results in (A). +, weak; ++, moderate; +++, strong; -, no localization. See Supplementary Figure 1 for expression in a wild-type (*Nin*^*+*^) background. See Supplementary Figure 2A for fluorescence intensity profiles.

Nin^1-1091^ localized to the nuclear surface in fat body cells (Figure 2 and Supplementary Figures 1A-B), mirroring antibody staining of endogenous Nin (Zheng *et al*., 2020). Localization was most prominent at the nuclear surface with a weaker signal present inside the nucleus (see fluorescence intensity profiles, Supplementary Figure 2). Perinuclear localization in punctate aggregates was variably present, which appeared to be due to overexpression as it was more prominent with two copies of the transgene than with one (Supplementary Figure 3A).

**Figure 3.**
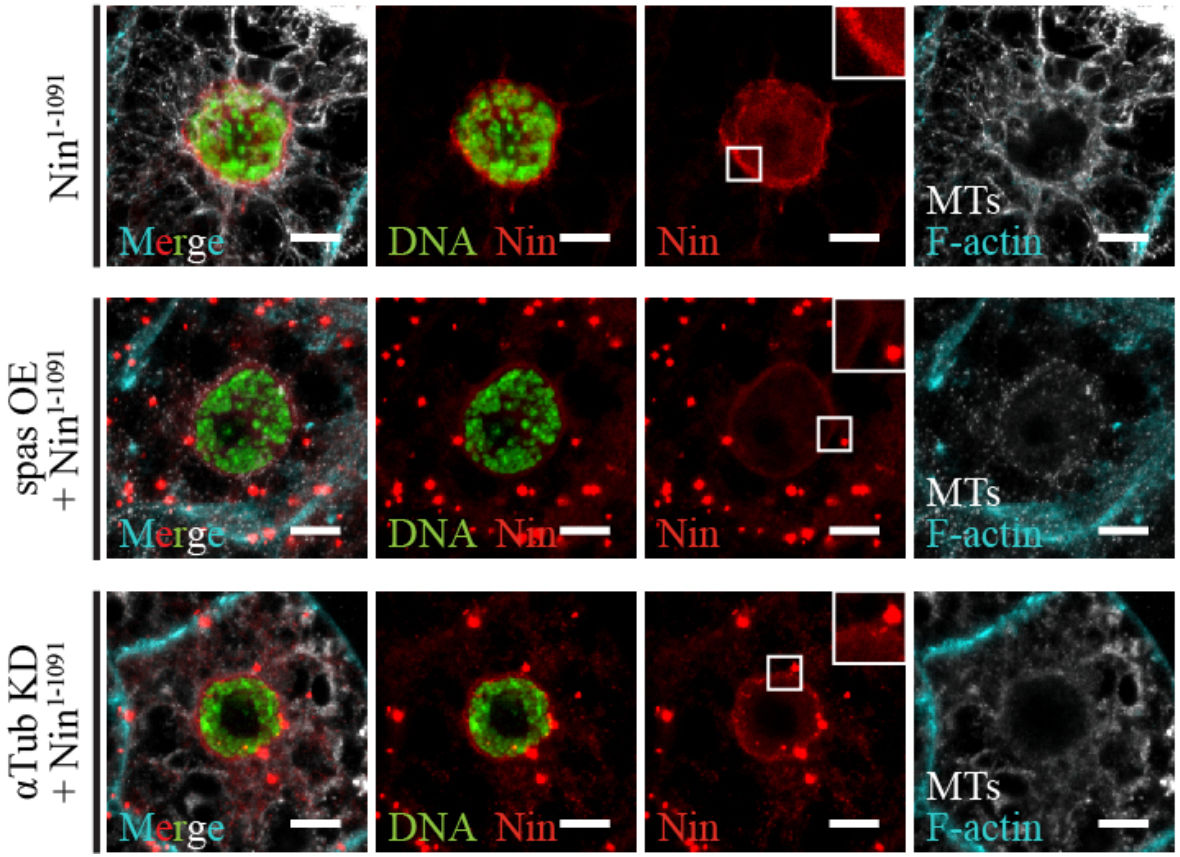
Localization of Ninein to the fat body ncMTOC is partially dependent upon microtubules. IF staining of larval fat body cells with DNA (DAPI, green), Nin^1-1091^ (TagRFP fluorescence, red), F-actin (488 Phalloidin, cyan), and microtubules (YL1/2, white). Insets show an enlarged view of the nuclear surface. Nin^1-1091^ localizes to the perinuclear MTOC and forms aggregates when microtubules are disrupted by the overexpression (OE) of *CFP-spastin* or by the RNAi-mediated knock-down (KD) of *α-tubulin*. See Supplementary Figure 4A for fluorescence intensity profiles.

Intranuclear signal also increased with two copies of Nin^1-1091^ (Supplementary Figures 3A). Expression of two Nin transgenes that express at higher levels, Eos-Nin and Nin-GFP (Supplementary Figure 3B), resulted in large intranuclear and cytoplasmic punctate aggregates (Supplementary Figure 3C), indicating that these intranuclear and aggregate patterns correlate with a high dosage effect of Nin overexpression. High levels of ubiquitous Nin overexpression are lethal (Zheng *et al*., 2016) with a restricted toxicity in the muscle or fat body (Table 1). Levels of Nin-TagRFP-Myc overexpression used in this study, however, were not lethal in the fat body using SPARC-GAL4 for each of the transgenes shown in Figure 1.

**Table 1.** Ninein overexpression is lethal in the muscle and fat body.

| Expression tissue | Driver (GAL4) | F1 phenotype |
| --- | --- | --- |
| Nervous system | Elav | Viable |
|  | Appl | Viable |
| Salivary gland | Sgs3 | Viable |
|  | Fkh | Viable |
| Midgut | Esg | Viable |
|  | Myo31DF | Viable |
| Hemocyte | He | Viable |
|  | Hml | Viable |
| Muscle | Mef2 | LETHAL |
| Fat body | SPARC | LETHAL |

Expression of Nin-TagRFP-Myc constructs containing N- (Nin^1-242^, Nin^1-353^, and Nin^1-451^) or C-terminal (Nin^568-1091^ and Nin^Δ451-567^) domains localized to the nuclear surface (Figure 2). An N-terminal fragment of amino acids 1-100 localized predominantly inside the nucleus; however, longer N-terminal constructs extending up to amino acid 451 were primarily outside the nucleus (Figure 2 and Supplementary Figure 1). Interestingly, a central region of Nin (Nin^454-571^) also localized intranuclearly but, unlike the N-terminal 100 amino acids, appeared to drive all fragments containing it into the nucleus (Figure 2 and Supplementary Figure 1). We used two programs to identify a nuclear localization signal (NLS) in these regions (see Materials and Methods), but none were detected from these programs. While amino acids 454-567 target all fragments that contain it to the nucleus, this region is less efficient at targeting the full-length protein into the nucleus possibly due to the presence of a competing domain(s) that localize Nin to the nuclear surface. Large fragments that lack this domain, including Nin^Δ451-567^, were excluded from the nucleus (Figure 2 and Supplementary Figure 1). These data indicate that at least two domains of Nin contribute to its localization to the nuclear surface while amino acids 454-567 drive the protein into the nucleus and amino acids 1-100 can passively localize there.

### MT-dependent and -independent modes of Nin localization to the MTOC

Because Nin binds MTs and is a MT anchor (Mogensen *et al*., 2000; Abal *et al*., 2002; Delgehyr *et al*., 2005; Kowanda *et al*., 2016), we next sought to determine whether MTs play a role in Nin localization at the nuclear surface. Disrupting MTs by overexpressing the MT-severing enzyme *spastin* or by knocking down the MT component *α-tubulin* by RNAi reduced, but did not eliminate, perinuclear localization of Nin^1-1091^ while increasing the incidence of punctae formation (Figure 3; see fluorescence intensity profiles, Supplementary Figure 4A). Thus, there appears to be MT-dependent and -independent modes of Nin localization to the nuclear surface.

**Figure 4.**
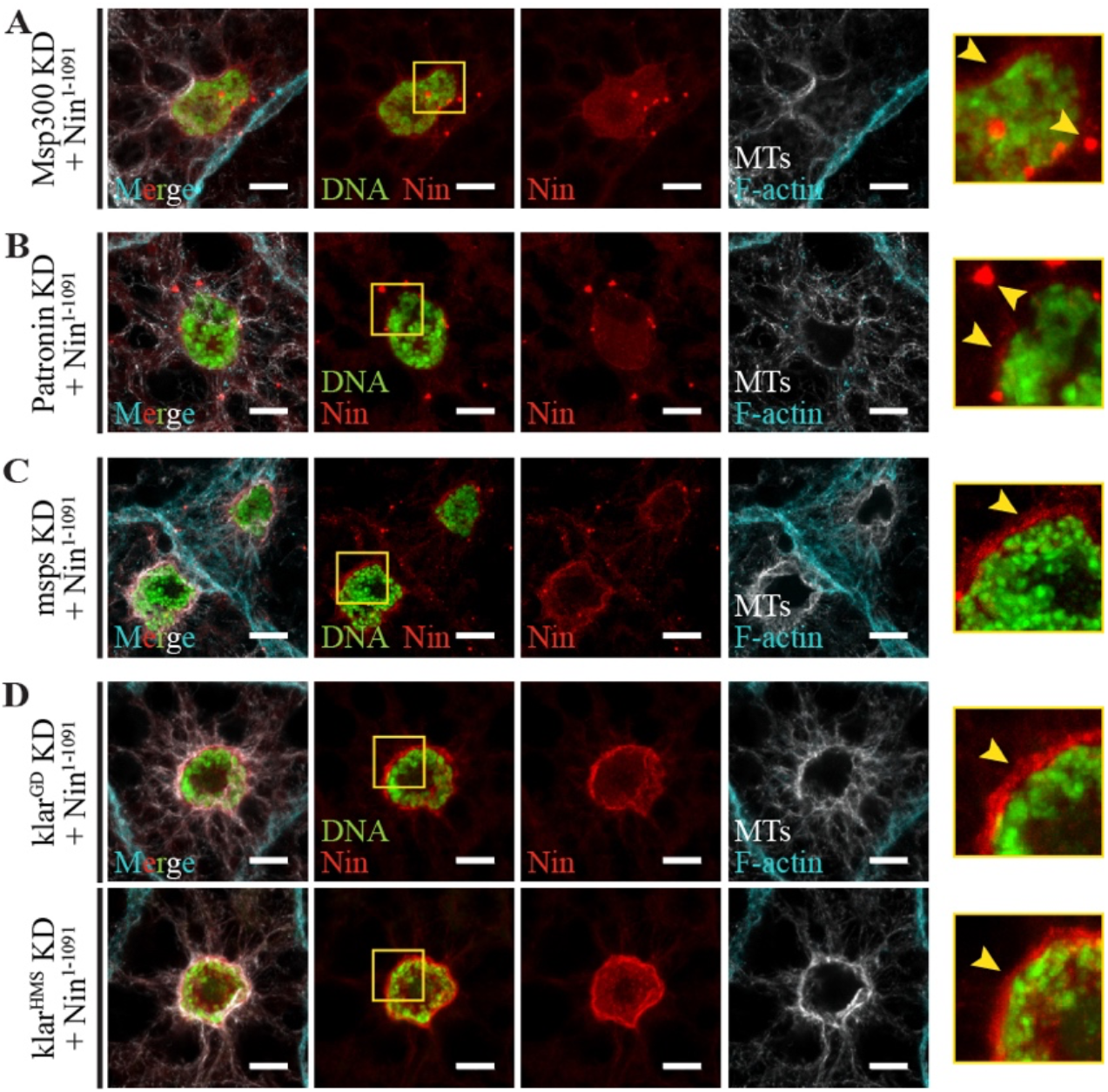
Fat body MTOC proteins Msp300 and Patronin are necessary for perinuclear Ninein localization. IF staining of larval fat body cells expressing Nin^1-1091^ using SPARC-GAL4 in a wild-type background with RNAi knock-down (KD) of the genes indicated. DNA (DAPI, green), Nin^1-1091^ (TagRFP fluorescence, red), F-actin (488 Phalloidin, cyan), and microtubules (YL1/2, white). Right panels show an enlarged view of the nuclear surface. (A-B) Knock-down of *Msp300* or *Patronin* reduces Nin localization to the nuclear surface and increases intranuclear localization of Nin and the formation of cytoplasmic aggregates. (C) Knock-down of *msps* does not affect Nin localization. (D) Knock-down of *klarsicht* (*klar*) using either of two RNAi lines (GD9271 above, HMS01612 below) does not alter Nin’s perinuclear localization. See Supplementary Figure 4B for fluorescence intensity profiles.

We next sought to determine which proteins anchor or recruit Nin to the nuclear surface. Our previous work showed that the Linker of Nucleoskeleton and Cytoskeleton (LINC) complex protein Msp300/Nesprin was a major structural component of the fat body MTOC required to recruit msps and Patronin to the nuclear surface to organize and assemble MTs there (Zheng *et al*., 2020). Knockdown of *Msp300* in the fat body disrupted nuclear positioning due to MT disruption (Zheng *et al*., 2020). Loss of *Msp300* (Figure 4A) or *Patronin* (Figure 4B) reduced perinuclear Nin and increased its intranuclear localization and cytoplasmic aggregate accumulation (arrowheads; see fluorescence intensity profiles, Supplementary Figure 4B).

However, we were unable to detect an effect on localization of Nin to the nuclear surface when *msps* (Figure 4C) or the other *Drosophila* Nesprin *klarsicht* (*klar*, Figure 4D) was knocked down using either of two RNAi lines (Figure 4D, arrowheads; see fluorescence intensity profiles, Supplementary Figure 4B). Therefore, Nin localization to the nuclear surface relies partly on MTs, but also on a MT-independent anchor that may involve Msp300 and Patronin. But whether these interactions are direct or indirect remains to be determined.

### Ensconsin but not dynein is required for Nin localization to the MTOC

We next analyzed whether Nin requires dynein and/or ens for proper localization. Loss of *Dynein heavy chain* (*Dhc*) or *Dynein light intermediate chain* (*Dlic*), a direct partner of NIN (Celestino *et al*., 2019; Lee *et al*., 2020), did not alter MT organization (Zheng *et al*., 2020) nor did it reduce perinuclear Nin localization (Figure 5A). Likewise, inactivation of dynactin via the overexpression of *DCTN2-p50*/*Dynamitin* did not affect MTs or prohibit Nin from localizing to the MTOC (Figure 5A; see fluorescence intensity profiles, Supplementary Figure 4C). Loss of *ens*, on the other hand, disrupted Nin localization to the nuclear surface resulting in perinuclear Nin aggregate formation (Figure 5B; see fluorescence intensity profiles, Supplementary Figure 4C).

**Figure 5.**
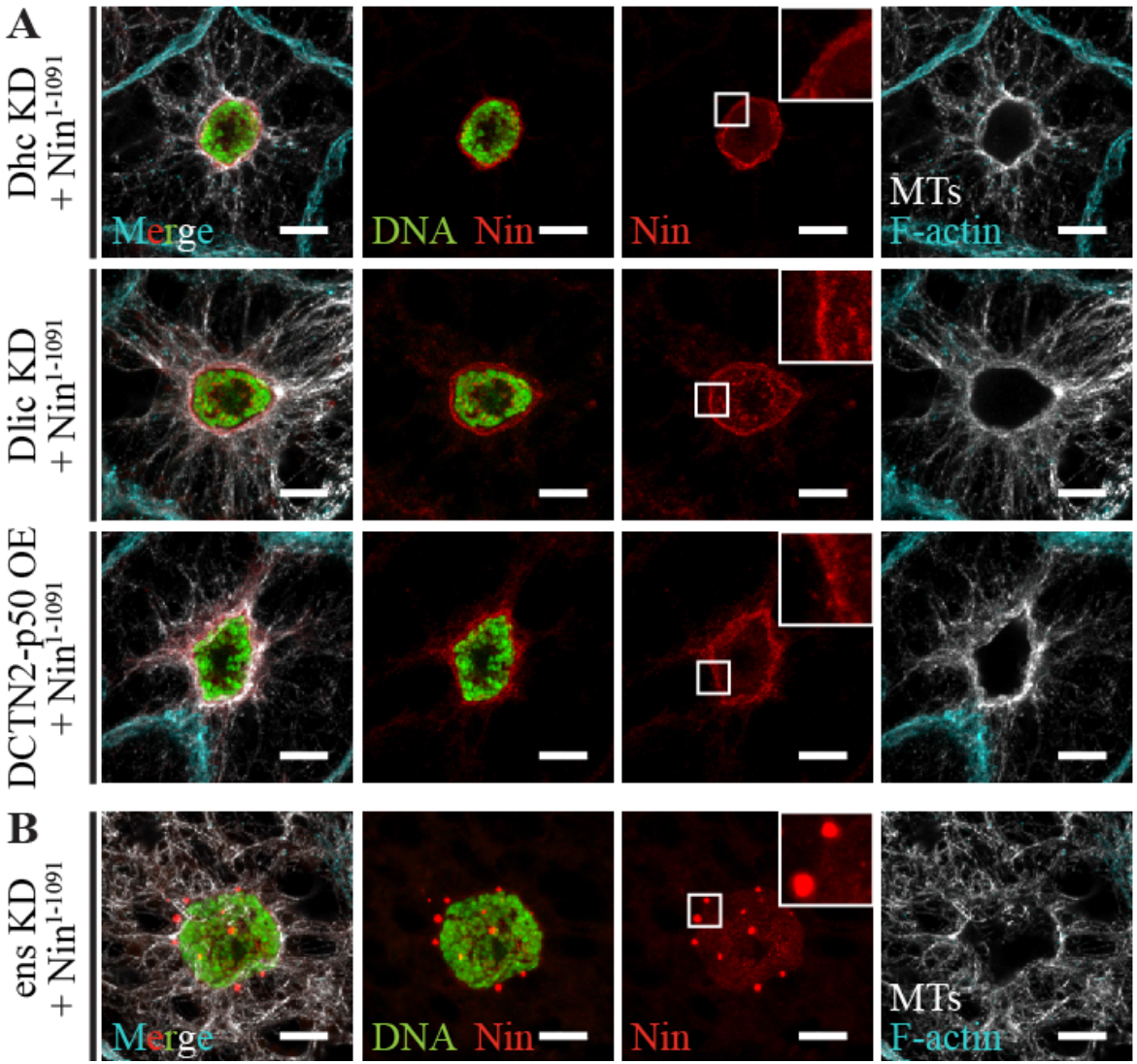
Ninein localization is dependent upon ens but not dynein. IF staining of larval fat body cells expressing Nin^1-1091^ using SPARC-GAL4 in a wild-type background with RNAi knock-down (KD) of the genes indicated. DNA (DAPI, green), Nin^1-1091^ (TagRFP fluorescence, red), F-actin (488 Phalloidin, cyan), and microtubules (YL1/2, white). Insets show an enlarged view of the nuclear surface. (A) Knock-down (KD) of *Dynein heavy chain* (*Dhc*) or *Dynein light intermediate chain* (*Dlic*) or inactivation of the dynein motor with *DCTN2-p50/Dynamitin* overexpression (OE) does not affect Nin localization to the MTOC. (B) Loss of *ensconsin* (*ens*) perturbs Nin localization, causing it to aggregate near the nucleus. See Supplementary Figure 4C for fluorescence intensity profiles.

### Ninein interacts with ensconsin and Dynein light intermediate chain

We next aimed to determine how Nin coordinates MT organization and if its partners play a role by first confirming whether Nin associates with ens and/or dynein in fat body cells. We overexpressed Nin-GFP in fat body cells using SPARC-GAL4, treated lysates with nocodazole on ice to depolymerize MTs, and pulled down Nin-GFP using GFP nanobody beads. We detected an association with endogenous ens (Figure 6A). Using the Nin truncation constructs (Figure 1), we further mapped the interaction domain by overexpressing ens-GFP together with various Nin constructs in fat bodies, pulling down ens-GFP, and probing for the Myc tag on Nin fragments. The C-terminus of Nin (Nin^568-1091^ and Nin^550-924^) co-immunoprecipitated with ens (Figure 6B, green arrows), but the N-terminus (Nin^1-571^) did not (Figure 6B, red arrow). We infer from the co-IP data the minimum interaction domain between Nin and ens to be within amino acids 572-924 of Nin (Figure 6B, see also Figure 7).

**Figure 6.**
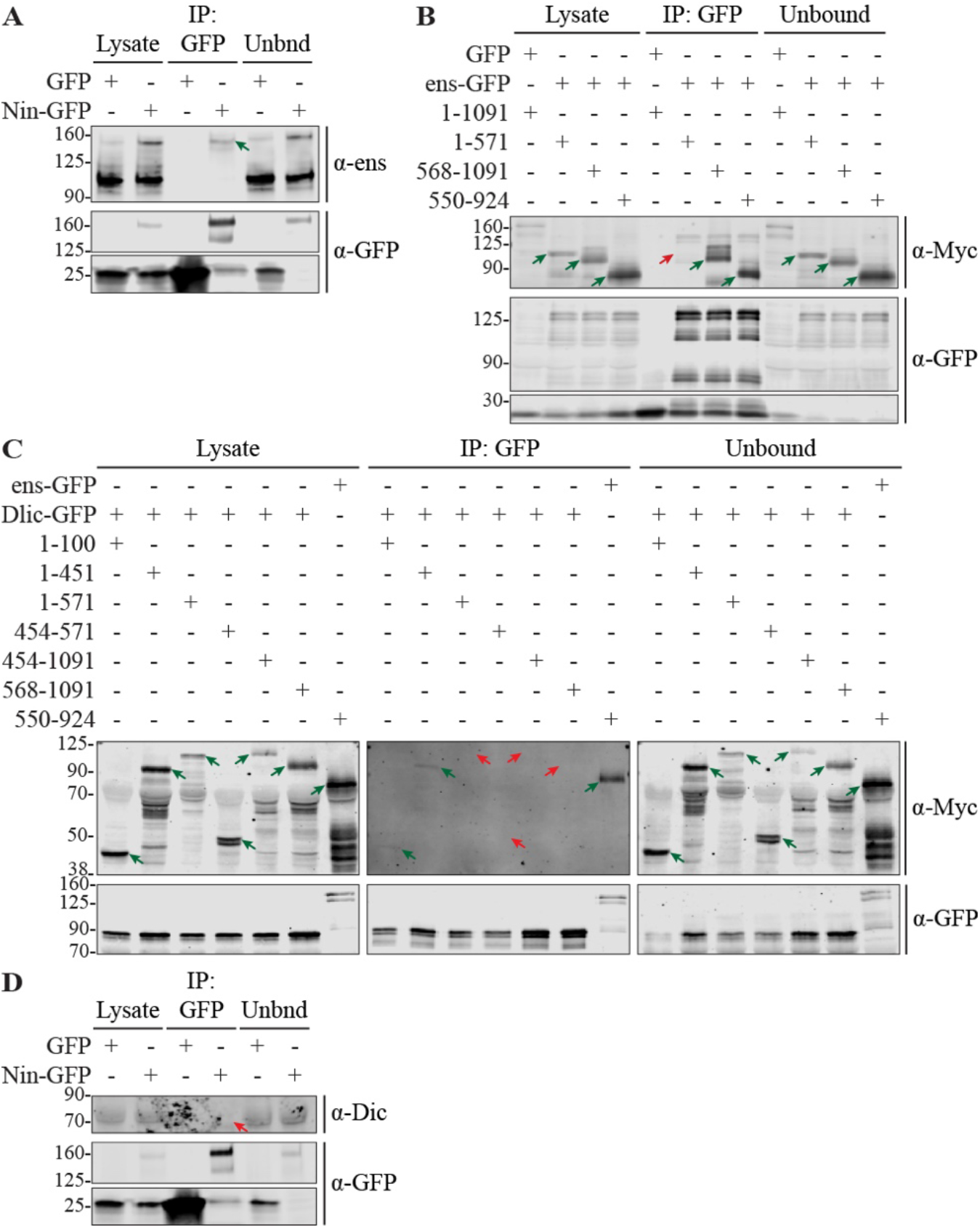
Mapping of Ninein’s ens- and Dlic-interaction domains. Western blot analysis of GFP immunoprecipitates from whole larval lysates expressing the indicated proteins in fat bodies. Unbnd = Unbound fraction in lysate after IP. Green arrows indicate a positive co-IP, red arrows a negative. (A) co-IP of ens with Nin-GFP. Membrane was probed for endogenous ens (∼150 and 100 kDa) and GFP. Endogenous ens (∼150 kDa) co-immunoprecipitated with Nin. (B) co-IP of Nin-TagRFP-Myc fragments with ens-GFP. Membrane was probed for GFP (ens-GFP; ∼180 kDa) and Myc (Nin; various sizes). (C) co-IP of Nin-TagRFP-Myc fragments with Dlic-GFP. Membrane was probed for GFP (Dlic-GFP; ∼81 kDa) and Myc (Nin; various sizes). Dlic co-immunoprecipitated with the N-terminal 100 amino acids of Nin. Nin^550-924^ + ens-GFP was used as a positive control. (D) Overexpression of Nin-GTP in fat bodies. Membrane was probed for endogenus Dynein intermediate chain (Dic; 74 kDa) and GTP. Dic co-IP was not detected (red arrow).

**Figure 7.**
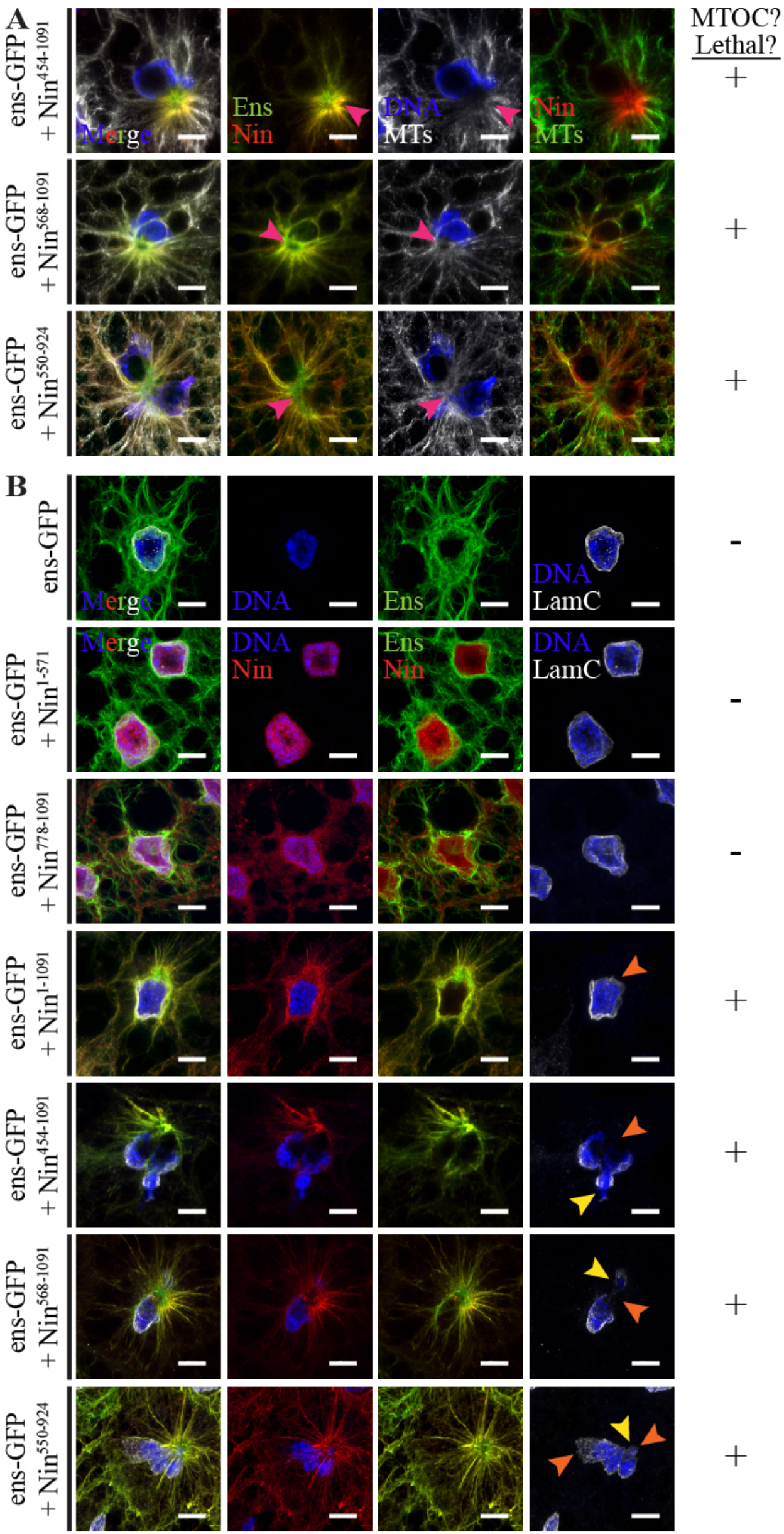
Ninein and ensconsin synergize to form an ectopic MTOC that disrupts nuclear morphology. DNA (DAPI, blue), Nin^1-1091^ (TagRFP fluorescence, red), Ens (GFP fluorescence, green), and microtubules (YL1/2, white) or LaminC (LC28.26, white). (A) Co-overexpression of ens and fragments that encompass amino acids 572-777 of Nin alters microtubule organization, forming a robust juxtanuclear MTOC (pink arrowheads) associated with late pupal lethality. (B) LamC staining labels the inner nuclear membrane and is reduced where Nin-ens ectopic MTOCs are formed (orange arrowheads). Additionally, nuclear DNA spills into the cytoplasm (yellow arrowheads).

Using a similar approach, we mapped the domain that binds dynein. In agreement with previous work that identified NIN as an activator of dynein (Redwine *et al*., 2017; Reck-Peterson *et al*., 2018) that binds directly to DLIC1 and DLIC2 via its N-terminal 87 amino acid EF hand domains (Celestino *et al*., 2019; Lee *et al*., 2020), the N-terminus of *Drosophila* Nin (Nin^1-100^ and Nin^1-451^) co-immunoprecipitated with Dlic-GFP (Figure 6C, green arrows) while central (Nin^454-571^) and C-terminal Nin fragments (Nin^454-1091^ and Nin^568-1091^) did not (Figure 6C, red arrow). A longer N-terminal fragment, Nin^1-571^, was not detected in the co-IP with Dlic-GFP as expected possibly due to lower expression levels. We were unable to detect an association between Dynein intermediate chain (Dic) and Nin (Figure 6D, red arrow), indicating a specific interaction between Nin and Dlic. A summary of the mapping of ens and Dlic binding domains to Nin is shown in Figure 8.

**Figure 8.**
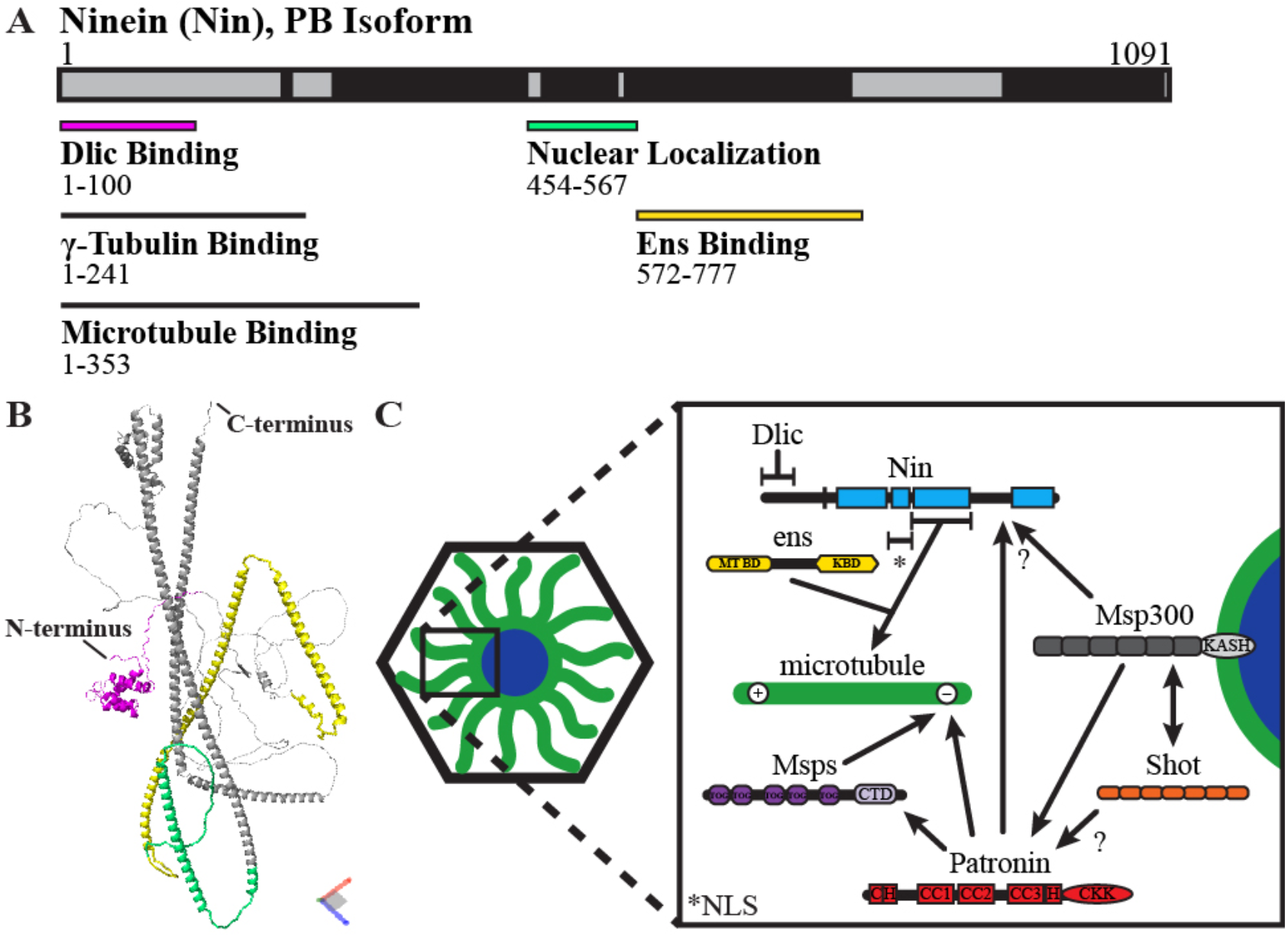
Summary of Ninein domains and model depicting its cooperative role with ensconsin at the MTOC. (A) Summary of Nin domains mapped from this work including the Dlic and ens binding domains and the nuclear localization domain. (B) Location of these domains on the predicted 3D structure of Nin-PB. Color coding matches that in (A). (C) Model for the proposed roles of Nin and ens at the fat body perinuclear MTOC, building on the model proposed recently (Zheng *et al*., 2020). KASH = Klarsicht, ANC-1, Syne Homology domain. CTD = C-Terminal Domain. CKK = CAMSAP1, KIAA1078, KIAA1543 domain. MT BD = microtubule binding domain. KBD = Kinesin binding domain.

### Ninein synergizes with ensconsin to promote MT assembly

Our next goal was to evaluate whether Nin’s interaction with dynein or ens was sufficient for MTOC formation. Ens and dynein heavy chain co-localize with MTs (Supplementary Figure 5A).

Surprisingly, we found that co-overexpression of constructs containing Nin’s C-terminus (Nin^454-1091^, Nin^568-1091^, and Nin^550-924^) with ens-GFP produced robust juxtanuclear MTOCs that superseded the perinuclear MTOC (Figure 7A). Expression of Nin or ens alone did not induce an ectopic MTOC, whereas together they synergize to organize MTs. These robust, ectopic MTOCs disrupted nuclear integrity as evidenced by alterations to the pattern of LaminC (LamC), a nuclear lamin nucleoskeleton component (Figure 7B, orange arrowheads). LamC staining was diminished at the nucleus adjacent to the positioning of the ectopic MTOC (Figure 7B, orange arrowheads), and chromosomal material appears to have spilled into the cytoplasm (Figure 7B, yellow arrowheads). We infer from these results that amino acids 572-777 of Nin are sufficient for the formation of an MTOC in cooperation with ens. This is consistent and overlaps with the ens-interacting domain identified from the co-IP experiments (see Figure 6B). Furthermore, this synergy is unique to the Nin-ens partnership, as overexpression of Nin^1-1091^ together with either Dlic or Dhc did not visibly alter MT organization (Supplementary Figure 5B).

## Discussion

In this study we generated a set of transgenic Nin deletion constructs in *Drosophila* that enabled us to map Nin’s localization, partner-binding, and MT-regulating domains *in vivo* in the larval fat body. From these data we identify domains of Nin that contribute to its localization inside the nucleus and to the MTOC on the nuclear surface, confirm the homologous N-terminal domain that associates with Dlic, and map the domain that associates with ens (Figure 8). From co-expression assays, we found a synergistic interaction between ens and Nin’s ens-binding domain that is sufficient to generate a robust ectopic MTOC.

Full-length tagged Nin (Nin^1-1091^) localizes primarily to the nuclear surface while lower levels are detected inside the nucleus. We further determined that a 114-amino acid domain at amino acids 454-567 is sufficient to target Nin into the nucleus (Figure 8A-B, green). All sub-fragments of Nin containing this domain showed significant localization within the nucleus; however, the full-length Nin protein had relatively low levels inside the nucleus possibly due to the presence of competing domains in the N- and C-terminal regions that target Nin to the nuclear surface.

Human DLIC1 was shown to bind directly to NIN via a pair of Ca++-independent EF hand domains at the N-terminal 87 amino acids of NIN (Lee *et al*., 2020), a region that is highly conserved between human and *Drosophila* nineins (Zheng *et al*., 2016). We confirm that *Drosophila* Nin also associates with Dlic and have mapped this interaction to the N-terminal end of Nin (Figure 8A-B, pink). This domain also localizes Nin to the nucleus where Dlic-GFP is localized, but longer N-terminal fragments containing this domain do not localize there (unless they also contain amino acids 451-567). The expected molecular weight of this domain with the engineered tag is ∼44 kDa, which is within the upper limit (30-60 kDa) of passive diffusion between nucleus and cytoplasm (Keminer and Peters, 1999; Wang and Brattain, 2007; Timney *et al*., 2016). This could explain why longer fragments such as Nin^1-242^ and Nin^1-353^ do not prominently localize inside the nucleus but instead are primarily localized to the perinuclear MTOC.

When Dlic-GFP was overexpressed in fat body cells, it localized in the nucleus. This was unexpected as Dlic’s role as a subunit of the dynein motor places its function with MTs in the cytoplasm. Other studies have used this and other tagged UAS-Dlic transgenes in tissues other than the fat body; however, their nuclear localization was not detected or reported (Pandey *et al*., 2007; Emre *et al*., 2011; Wainman *et al*., 2012; Baumbach *et al*., 2015; Inaba *et al*., 2015).

Notably, Dlic-GFP overexpression drove Nin into the nucleus. This nuclear localization dynamic between Dlic and Nin is enigmatic but does not seem to overtly impact MT organization. Furthermore, loss of either Dhc or Dlic or inactivation of dynactin does not affect Nin localization to the perinuclear MTOC or MT organization in fat body cells. Although the interaction between Nin and Dlic may be important in other cellular processes, our assays did not reveal a role for it in MT organization in the fat body.

N- and C-terminal Nin-TagRFP-Myc constructs localize to the MTOC. The contribution of multiple domains of Nin conferring localization to the MTOC may point to its interactions with several distinct partners including itself, MTs, ens, etc. that anchor Nin at the MTOC. The fat body nuclear surface has prominent circumferential MTs (Zheng *et al*., 2020) and recruitment of full-length Nin is largely, but not completely, dependent on these MTs. Depolymerizing MTs may release a pool of Nin that interacts with MTs so that Nin is free to aggregate, leading to a phenotype similar to high levels of Nin overexpression. How Nin anchors to the MTOC independent of MTs has yet to be determined but may involve Msp300, Patronin, and/or ens.

From the co-IP data, we identified a Nin-ens interaction domain between amino acids 572-924 of Nin. Through staining, we further refined this domain to amino acids 572-777 (Figure 8A-B, yellow). We further demonstrated that, when overexpressed, Nin and ens function synergistically to form an ectopic ncMTOC. When Nin constructs that include amino acids 572-777 are overexpressed with ens, robust ectopic MTOCs are formed adjacent to the nucleus. Interestingly, Nin’s interaction with ens is sufficient to overcome Nin’s nuclear localization (compare Nin^454-1091^ Figure 7A-B to Figure 2A). In contrast to the developing muscle where multiple foci formed from Nin-ens overexpression (Rosen *et al*., 2019), there is only one Nin-ens MTOC per fat body cell. Overexpression of Nin-TagRFP-Myc fragments or ens-GFP alone in the fat body produced no overt effects and flies were viable, whereas co-overexpression of Nin’s ens-binding domain together with ens-GFP was late pupal lethal. In contrast with these synergistic effects of Nin and ens expression in the fat body, similar overexpression experiments in the developing muscle showed that Nin overexpression was lethal but was suppressed by co-overexpression of ens (Rosen *et al*., 2019). Whether the MTOC that Nin-ens generates involves other factors remains to be determined. It likely does not require *γ*-tubulin because *γ*-tubulin is expressed at very low levels in the fat body and was not required for MT assembly at the primary fat body MTOC (Zheng *et al*., 2020).

In the fat body, Nin-ens ectopic MTOCs were positioned near the nucleus, disrupted the nuclear envelope, altered nuclear morphology, and resulted in the leakage of chromosomal material into the cytoplasm. One function of the fat body MTOC under normal conditions may be to ensure proper nuclear morphology. Normally, MTs in the fat body are organized both in circumferential bundles surrounding and with their minus ends anchored at the nuclear surface (Zheng *et al*., 2020). With the generation of Nin-ens ectopic MTOCs, the nuclear morphology changes could be the result of the forces generated by imposing MTs emanating from the newly established ectopic MTOC. The disruption of LaminC patterning at the nuclear periphery is consistent with the fragility of these nuclei, as mutations in lamins can produce a similar phenotype (Davidson and Lammerding, 2014). Moreover, impinging MTs can impact nuclear morphology, a phenomenon also correlated with reduction of lamin signal (Biedzinski *et al*., 2020; Heffler *et al*., 2020). This effect may explain the nuclear morphology changes and loss of structural integrity caused by the fat body Nin-ens ectopic MTOC. Presumably the lethality associated with Nin and ens co-overexpression is due to these nuclear disruptions.

Altogether, our findings reveal novel features of domains required for Nin’s localization to the fat body MTOC on the nuclear surface, localization inside the nucleus, its interactions with dynein and ens, and how those interactions impact localization of Nin and MT organization in fat body cells. Furthermore, we identify a synergistic effect between Nin and ens capable of coordinating an MTOC, implying a mechanistic connection between this partnership normally at the fat body perinuclear and possibly at other MTOCs. It will be interesting to discover whether human NIN has a cooperative role with MAP7 (ens ortholog) in organizing MTs and whether this connection has disease relevance.

## Materials and Methods

### Generation of UAS-Nin-TagRFP-Myc Transgenes

The Nin coding sequence (*Nin-RB* isoform) was amplified by polymerase chain reaction (PCR) from the LD21844 cDNA clone (RRID:DGRC_5314) using the primers 1.FWD and 1.REV. The PCR product which included the entire ORF was cloned into the pENTR/D-TOPO vector (ThermoFisher, Cat#K240020). Using Gateway LR cloning, the coding sequence was inserted into pBID-UASC-GRM (RRID:Addgene_35203, (Wang *et al*., 2012)), creating a C-terminal TagRFP-Myc-tagged Nin construct (pBID-UASC-Nin-GRM). A version of pBID-UASC-Nin-GRM was also generated to be RNAi resistant to the Nin[HMS23837] RNAi line (BDSC #62414). The primers used to create this plasmid introduced silent mutations (CA<u>A</u>GA<u>G</u>AT<u>TTCAAGT</u>CT<u>C</u>CA<u>G</u>, mutations bold and underlined) in the RNAi recognition motif (CAGGAAATCAGTTCACTGCAA). Either of these plasmids was used as a template for the construction of Nin constructs shown in Figure 1. PCR fragments were inserted into EcoR1- and Bsu36I-digested pBID-UASC-GRM plasmid using NEBuilder HiFi DNA Assembly (New England BioLabs Inc., Cat#E5520S). Primers were designed using SnapGene and purchased from Integrated DNA Technologies, Inc. Primers and sequences are shown below. Transgenes were then inserted at VK40(3R) attP docking site by ΦC31 integration by GenetiVision, Inc. and screened in our lab. Transgenics were selected by expression of mini-white and confirmed by Western blotting and *in vivo* expression.

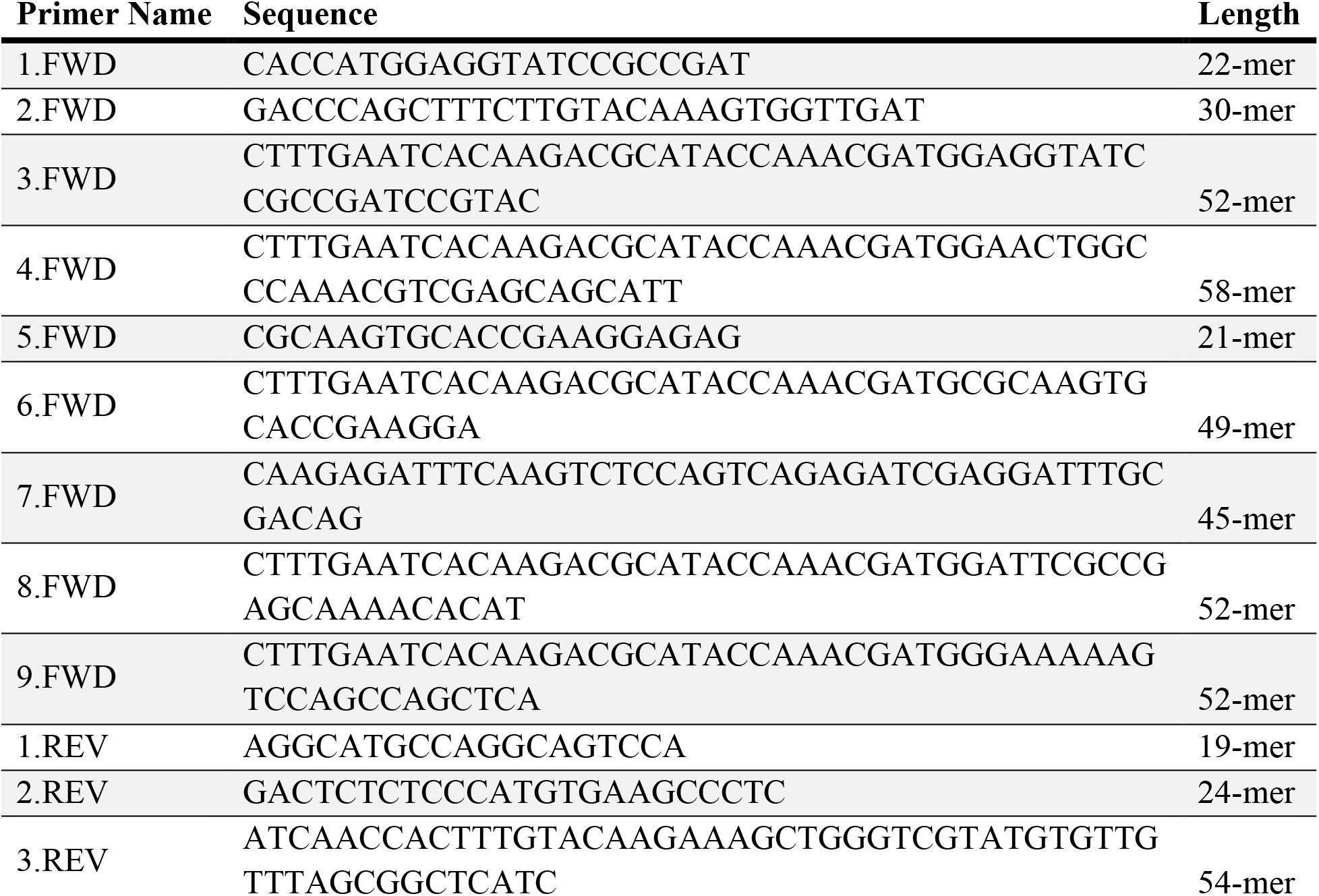
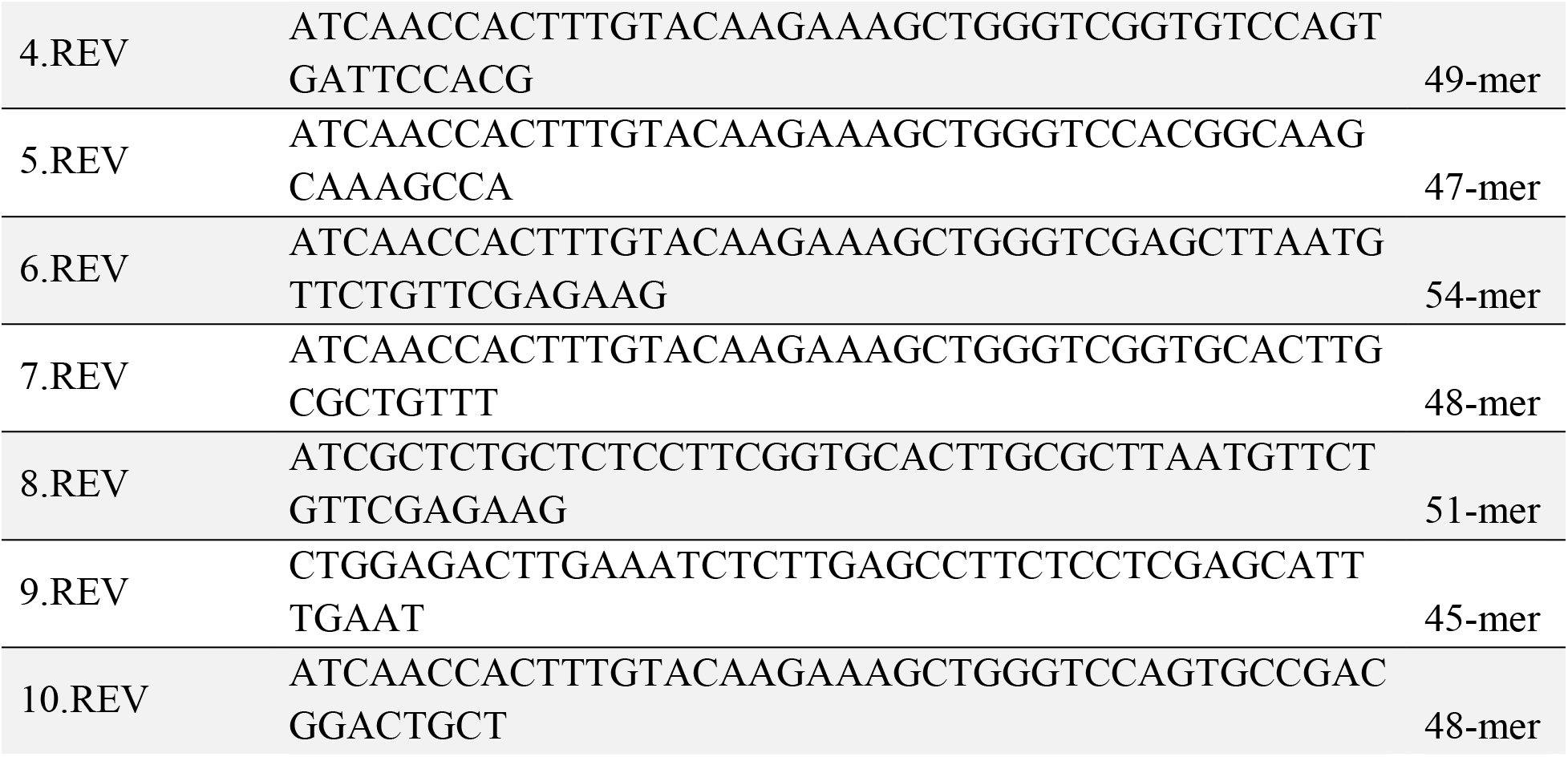

### Fly Stocks

Flies were maintained on standard food. Crosses were conducted at 29° unless otherwise stated. For crosses involving tub-GAL80^ts-7^, crosses were started at 25° and then moved to 29° on the third day. *w*^*1118*^ was used as wild-type.

Identifiers and sources for the fly strains used in this study can be found below (Bloomington *Drosophila* Stock Center = BDSC, Vienna *Drosophila* Resource Center = VDRC):

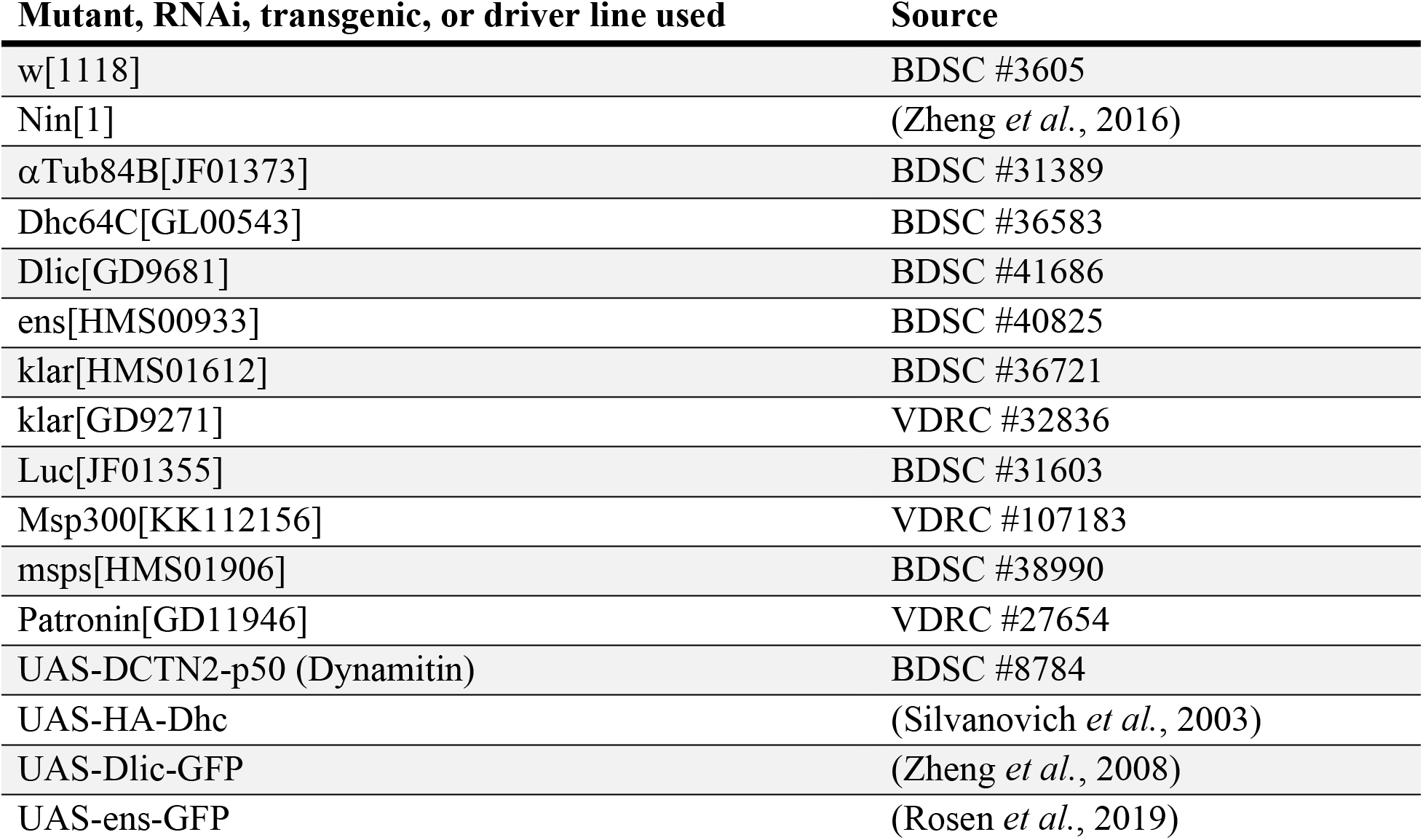
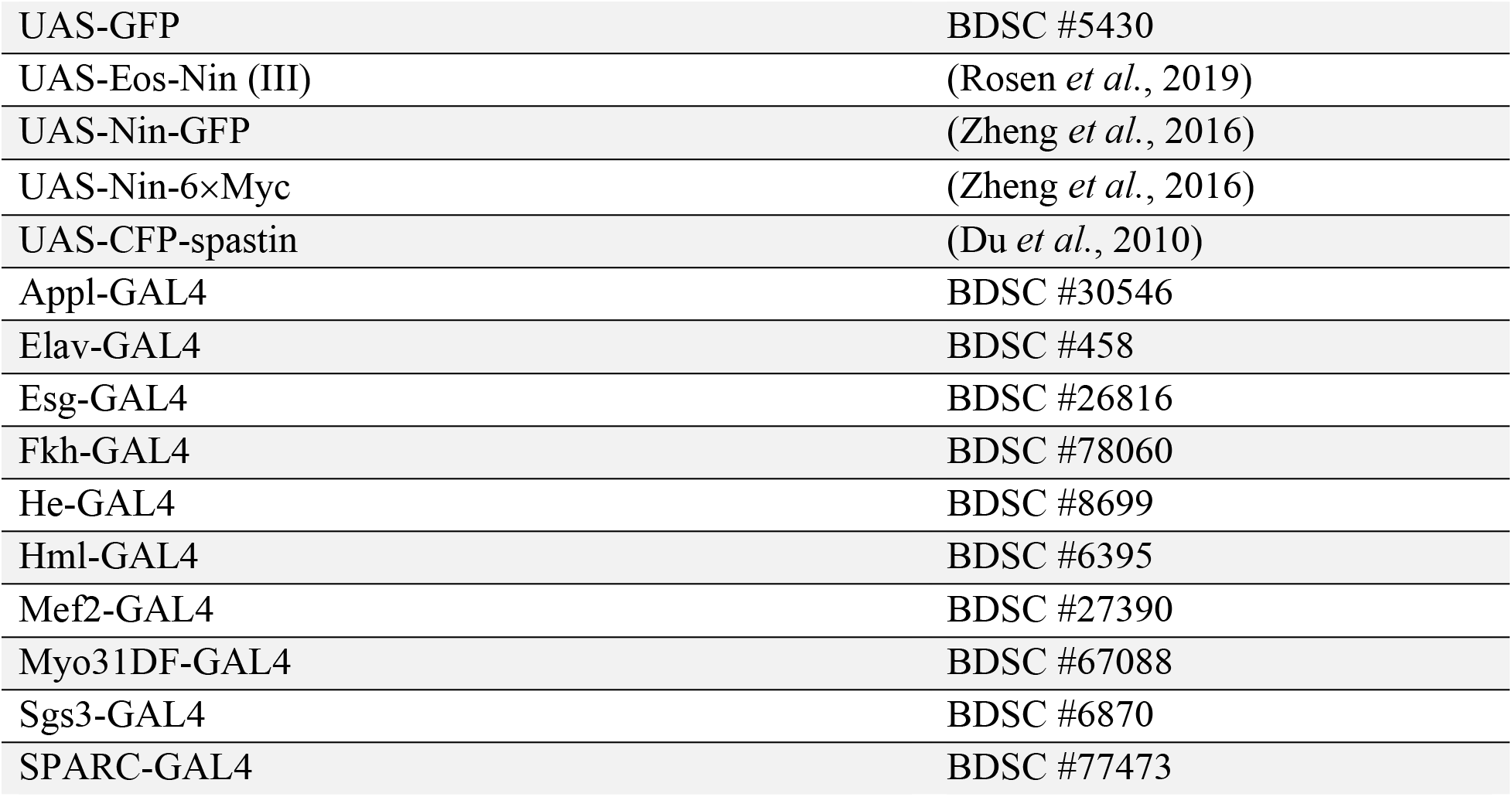

| Mutant, RNAi, transgenic, or driver line used | Source |
| --- | --- |
| w[1118] | BDSC #3605 |
| Nin[1] | (Zheng *et al.*, 2016) |
| αTub84B[JF01373] | BDSC #31389 |
| Dhc64C[GL00543] | BDSC #36583 |
| Dlic[GD9681] | BDSC #41686 |
| ens[HMS00933] | BDSC #40825 |
| klar[HMS01612] | BDSC #36721 |
| klar[GD9271] | VDRC #32836 |
| Luc[JF01355] | BDSC #31603 |
| Msp300[KK112156] | VDRC #107183 |
| msps[HMS01906] | BDSC #38990 |
| Patronin[GD11946] | VDRC #27654 |
| UAS-DCTN2-p50 (Dynamitin) | BDSC #8784 |
| UAS-HA-Dhc | (Silvanovich *et al.*, 2003) |
| UAS-Dlic-GFP | (Zheng *et al.*, 2008) |
| UAS-ens-GFP | (Rosen *et al.*, 2019) |
| UAS-GFP | BDSC #5430 |
| UAS-Eos-Nin (III) | (Rosen *et al.*, 2019) |
| UAS-Nin-GFP | (Zheng *et al.*, 2016) |
| UAS-Nin-6×Myc | (Zheng *et al.*, 2016) |
| UAS-CFP-spastin | (Du *et al.*, 2010) |
| Appl-GAL4 | BDSC #30546 |
| Elav-GAL4 | BDSC #458 |
| Esg-GAL4 | BDSC #26816 |
| Fkh-GAL4 | BDSC #78060 |
| He-GAL4 | BDSC #8699 |
| Hml-GAL4 | BDSC #6395 |
| Mef2-GAL4 | BDSC #27390 |
| Myo31DF-GAL4 | BDSC #67088 |
| Sgs3-GAL4 | BDSC #6870 |
| SPARC-GAL4 | BDSC #77473 |

The following lines were generated for this study:

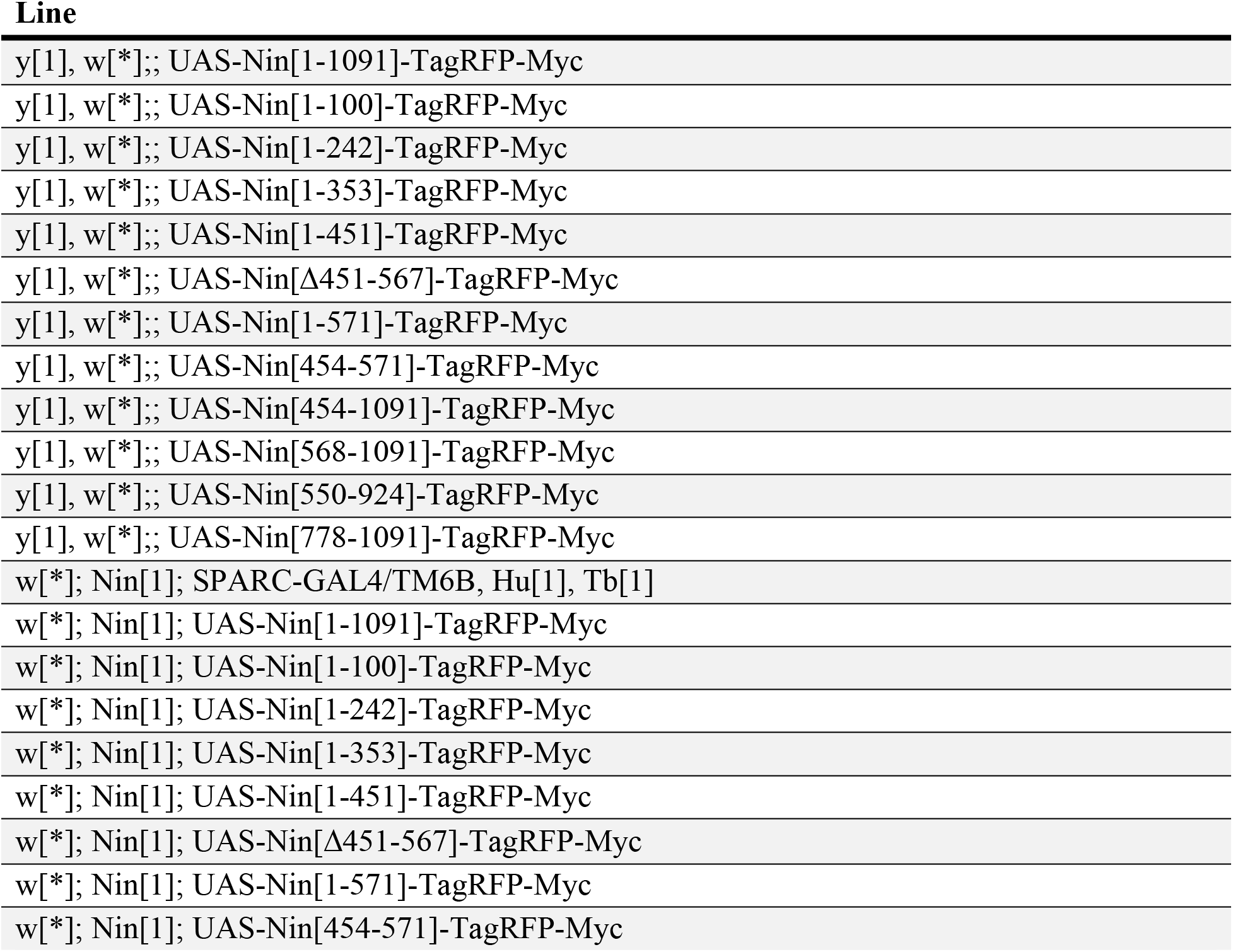
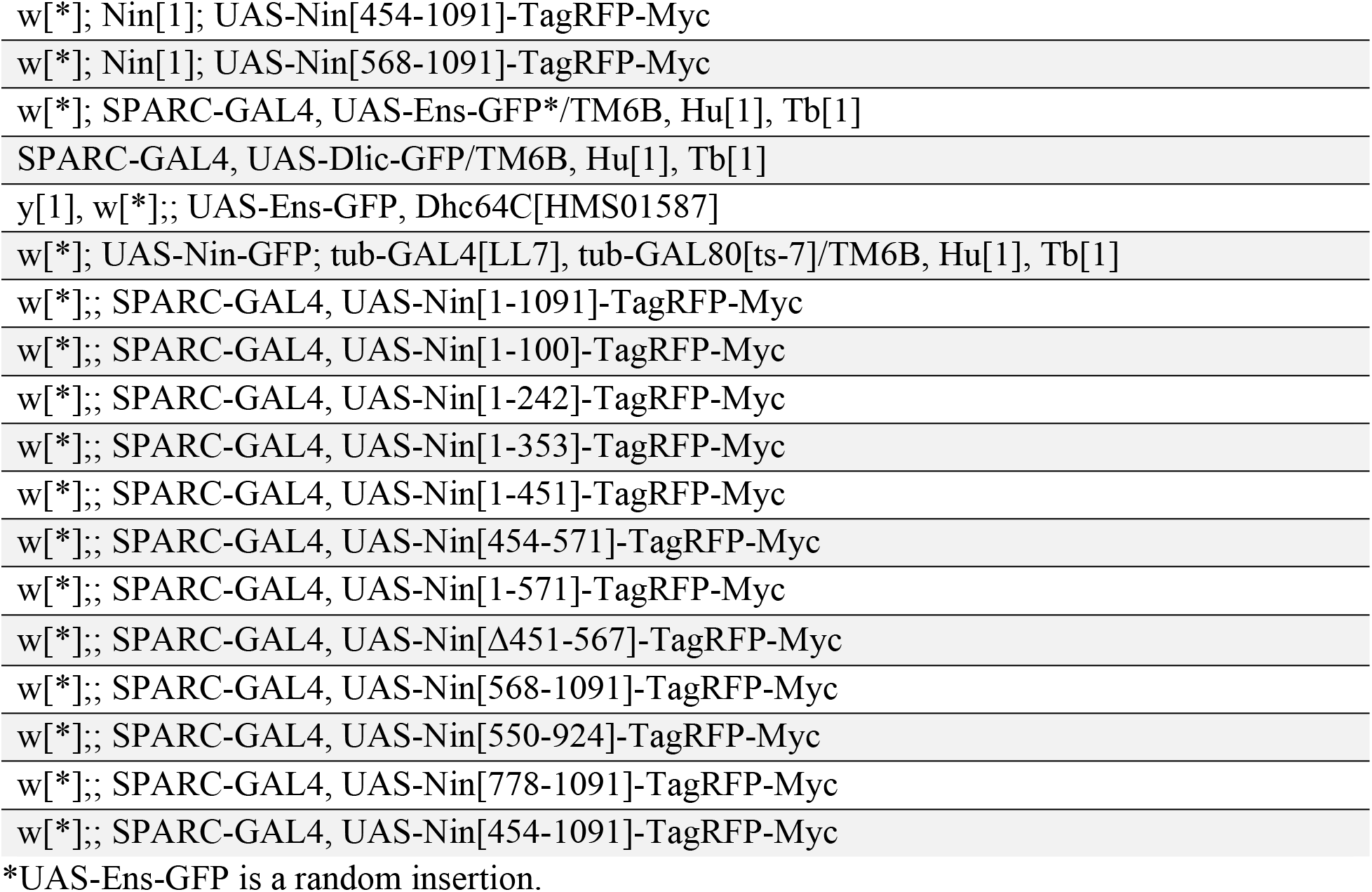

### Survey of Ninein Lethality

The indicated driver was crossed to UAS-Nin-GFP (Zheng *et al*., 2016). Progeny were screened for viability (i.e., eclosion from pupal case).

### Nuclear Localization Signal Database Search

The sequence of Nin PB from amino acids 1-100 and 454-567 were separately analyzed with the following programs to check for nuclear localization sequences: NLStradamus (Nguyen Ba *et al*., 2009) and Nuclear Localization Signal Predictor (NovoPro). Default settings were used for each program.

### Immunostaining and Imaging

Fat bodies were dissected from two wandering third instar larvae in 1X DPBS (Gibco, Ref#14080-055) and mounted on poly-Lysine treated (Kao and Megraw, 2004) slides (VWR, Cat#89085-339) in 12 *μ*L of 4% PFA (Spectrum, Cat #P1010). 2-4 slides were prepared for each condition. After ∼8 minutes in PFA, a siliconized 22×22 coverslip (VWR, Cat#48366-227) was applied, and the tissues were allowed to flatten for ∼1 minute before the slide was snap frozen in liquid nitrogen. Once frozen, the slide was removed from the liquid nitrogen and the coverslip was quickly pried off using a razor blade. The slide was then immediately placed into a Coplin jar filled with PBS, pH 7.3 (137 mM NaCl (OmniPur, Cat#7760), 2.7 mM KCl (OmniPur, Cat#7300), 8 mM Na2HPO4 (VWR, Cat#0404-500G), 1.4 mM KH2PO4 (Millipore, Cat#529568-250GM)) prior to application of primary antibodies.

Once all samples had been processed, slides were dried, and a hydrophobic ring was drawn around the tissue sample using a Liquid Blocker Super PAP Pen (Electron Microscopy Sciences, Cat#71312). The sample was incubated overnight at 4°C with a blocking solution (PBS, 0.5% BSA (Boehringer Mannheim Corp., Cat#100-021), 0.1% saponin (Sigma, Cat#S-2149)) in a dark humid chamber. The next day, slides were incubated with primary antibody either overnight at 4°C or for 4-6 hours at room temperature. Following incubation with primary antibodies, samples were washed 3×10 minutes with PBS. Samples were then incubated with secondary antibodies for 1.25 hours at room temperature. After secondary antibody incubation, samples were washed 3×5 minutes with PBS and then mounted in 15 *μ*L of mounting media (80% Glycerol (Alfa Aesar, Cat#36646); 0.1 M Tris•HCl (Calbiochem, Cat#648311), pH 8.8; 0.05% p-phenylenediamine (Sigma Aldrich, Cat#P-1519)). Slides were imaged using a Nikon A1R confocal microscope with an Apo TIRF 60X/1.49 oil objective using NIS-Elements AR 4.6 software.

### Antibodies and Stains

The following antibodies were used in this work: rat anti-α-tubulin (YL1/2; 1:1000 IF; 1:3333 immunoblotting (WB); Invitrogen, RRID:AB_2210201), mouse anti-Dhc (2C11-2; 1:50 IF; DSHB deposited by J. M. Scholey, RRID:AB_2091523, (Sharp *et al*., 2000)), mouse anti-Dic (MAB1618; 1:1000 WB; EMD Millipore, RRID:AB_1674698), rabbit anti-ensconsin (1:20 IF; 1:1000 WB; gift from Vladimir Gelfand, (Barlan *et al*., 2013)), rabbit anti-HA tag (C29F4; 1:1000 IF; Cell Signaling Technology, Cat# 3724, RRID:AB_1549585), chicken anti-GFP (1:5000 WB; Aves Labs, RRID:AB_2307313), rabbit anti-GFP (1:10000 WB; Invitrogen, RRID:AB_221570), mouse anti-LaminC (LC28.26; 1:100 IF; DSHB deposited by P. A. Fisher, RRID:AB_528339, (Riemer *et al*., 1995)), mouse anti-Myc tag (9B11; 1:5000 WB; Cell Signaling Technology, RRID:AB_331783), and guinea pig anti-Ninein (1:1000 WB; gift from Eric Lécuyer). The following stains were used for IF staining: DAPI (DNA; 1 *μ*g/mL; Sigma) and Phalloidin-iFluor 488 Conjugate (F-actin; 1:1000; Sigma).

Nin transgenes are inherently fluorescent due to the TagRFP tag and did not require additional staining. Likewise, Ens-GFP and Dlic-GFP did not require staining.

### Fluorescence Intensity Profile Measurements

Captured immunostaining images were opened in NIS-Elements AR 4.6 software > Measure > Intensity Profile. A single *z*-slice near the center of the image was selected for measurement and a ∼30 *μ*m line was drawn bisecting the nucleus. The fluorescence intensity profile was calculated automatically by the software.

### co-Immunoprecipitation

20-40 larvae were collected, washed in water, dried on a paper towel, and then frozen at -80°C until all samples and controls had identical numbers of larvae. Using a micro pestle (VWR International Pestle, Cat#47747-366) and mechanical tissue homogenizer (VWR International Pellet Mixer, Ref#47747-370), larvae were homogenized in 500 *μ*L of Lysis Buffer (10 mM Tris•HCl, pH 7.5; 150 mM NaCl; 0.5 mM EDTA (OmniPur, Cat#4005); 0.5% Nonidet P40 Substitute (VWR, Cat#E109-100ML)) containing protease inhibitors (1 mM 1,10-phenanthroline monohydrate (Sigma-Aldrich, Cat#P9375-5G); 0.5 mM PMSF (Sigma, Cat#EM-7110); 1 mM benzamidine hydrochloride hydrate (Sigma-Aldrich, Cat#B6506-5G); 1X protease inhibitor cocktail (Sigma-Aldrich, Cat#P8340-1ML)) and 0.4 mM nocodazole (Sigma, Cat#M1404-2MG). The lysates were incubated on ice for 30 minutes with light vortexing every 10 minutes. Then, lysates were centrifuged at 15,000*xg* for 6-7 minutes at 4°C and the cleared lysate was added to a pre-cooled tube containing 300 *μ*L of Dilution Buffer (10 mM Tris·HCl, pH 7.5; 150 mM NaCl; 0.5 mM EDTA) with protease inhibitors (identical to Lysis Buffer). This diluted lysate was then incubated with equilibrated GFP-Trap Magnetic beads (Chromotek, Cat#gtmak, RRID:AB_2631358) for 45 minutes-1 hour at 4°C with agitation. Following incubation, beads were washed 3×15 minutes with Dilution Buffer with protease inhibitors before being resuspended in 2X SDS-PAGE Loading Buffer (100 mM Tris·HCl, pH 6.8; 4% SDS (J.T. Baker, Cat#4095-02); 0.02% Bromophenol Blue (FisherBiotech, Cat#BP115-25); 20% Glycerol; 5% BME (OmniPur, Cat#6010)) for immediate analysis via Western blotting. For co-IP Western blots, input and unbound fractions represent ∼1% of the total lysate and IP fractions represent 45-50% of the total immunoprecipitate. To map the Dlic interaction domain, 100 larvae were collected and processed as above in 1250 *μ*L of Lysis Buffer. co-IP of any two proteins was repeated at least three times with the exception of Nin-GFP pull-down probing for endogenous DIC.

### Western Blotting and Analysis

To verify the Nin transgenes generated for this study, SPARC-GAL4 was used to drive expression of the transgenes in the fat body. Two larvae were collected, washed, and lysed using a micro pestle and mechanical tissue homogenizer in 40 μL of 2×SDS-PAGE Loading Buffer. After incubating at 95°C for 5 min, larval lysates were quickly spun down and 6 μL was loaded for SDS-PAGE gel electrophoresis on an 8% SDS-PAGE gel. The gel was transferred using a wet high intensity field transfer for 1 hour onto nitrocellulose membrane. The membrane was then blocked with 5% nonfat milk (Publix, Instant Nonfat Dry Milk) in TBS (0.5 M Tris·HCl, pH 7.5; 1.2 M NaCl) for 1 hour at room temperature or overnight at 4°C with rocking. This was followed by a brief wash in TBST (TBS with 0.1% Tween 20 (Fisher Bioreagents, Cat#BP337-100)) to rinse out the milk. The membrane was then probed with primary antibodies diluted in TBST for 1.25 hours at room temperature or overnight at 4°C with rocking. After washing with TBST 3×10 minutes, the membrane was incubated with secondary antibodies conjugated with IRDye-800CW or IRDye-680LT (1:20,000, LI-COR) for 1.25 hours at room temperature with rocking. Blots were scanned on an Odyssey CLx-2666 Infrared Imager (LI-COR Biosciences) using Image Studio v 5.2.5 software (LI-COR).

To measure the expression of various Nin transgenes in the fat body, fat bodies were dissected from three wandering third instar larvae. Genital discs remained, but all other tissues and glands were removed. Fat bodies were then homogenized by pipetting in 20 μL of 2×SDS-PAGE Loading Buffer and analyzed by Western blotting as above on a 10% SDS-PAGE gel.

## Abbreviations

co-IP: co-Immunoprecipitation
DCTN2-p50/Dynamitin: Dynactin 2, p50 subunit
Dhc: Dynein heavy chain
Dic: Dynein intermediate chain
Dlic: Dynein light intermediate chain
ens: ensconsin
MAP7: microtubule associated protein 7
IF: immunofluorescence
klar: klarsicht
LamC: LaminC
LINC: Linker of Nucleoskeleton and Cytoskelton
MT: microtubule
msps: mini spindles
Msp300: Muscle-specific protein 300 kDa
MTOC: microtubule-organizing center
ncMTOC: non-centrosomal MTOC
Nin: Ninein
Nlp: Ninein-like protein
NLS: nuclear localization signal/sequence
PCM: pericentriolar material
KD: RNAi knock-down
tub: tubulin
WB: immunoblotting/Western blot

## Acknowledgments

We thank Eric Lécuyer for antibodies to Nin, Vladimir Gelfand for antibodies to ensconsin, and all the contributors to the Developmental Studies Hybridoma Bank (DSHB) (see Materials and Methods) for providing antibodies used in this study. DSHB was created by the NICHD of the NIH and is maintained at The University of Iowa, Department of Biology, Iowa City, IA 52242. We thank Michael Welte for sharing his expertise on LINC complex proteins and functions, discussing the project during its infancy with us, and providing a critical reading of the manuscript. We thank Mary Baylies for UAS-Eos-Nin transgenic stocks, Tom Hays for UAS-HA-Dhc, Yuh-Nung Jan for UAS-Dlic-GFP, and Melissa Rolls for UAS-CFP-spastin. Many thanks to Bloomington Drosophila Stock Center, Vienna Drosophila Resource Center, and FlyBase for curating indispensable tools and resources. Thank you to Batory Foods for their generous donation of fly food reagents to support this work. We are also grateful to the members of the Megraw Lab for their countless hours of helpful discussion.

This work was supported by a Legacy Fellowship to Marisa Tillery from Florida State University and NIH grant R01GM139971 to Timothy Megraw.

## Author contributions

MT and TM designed the study and assembled the manuscript. MT performed the experiments with the exception of the Nin overexpression lethality screen performed by YZ. CZ constructed previously published Nin plasmids and provided experimental guidance and troubleshooting. All authors analyzed the data and approved the final version of the paper.

