## Supplementary Figures 1-5 for "Ninein domains required for its localization, association with partners dynein and ensconsin, and microtubule organization"

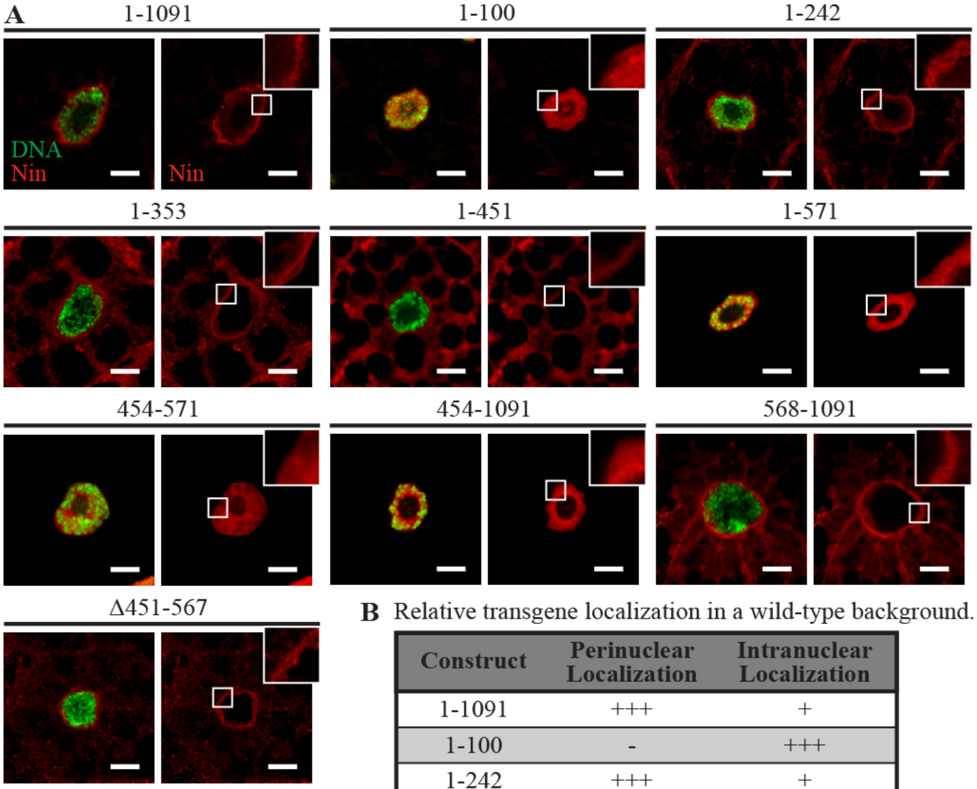


**B** Relative transgene localization in a wild-type background.

| Construct | Perinuclear Localization | Intranuclear Localization |
|---|---|---|
| 1-1091 | +++ | + |
| 1-100 | - | +++ |
| 1-242 | +++ | + |
| 1-353 | ++ | - |
| 1-451 | + | - |
| 1-571 | - | +++ |
| 454-571 | - | +++ |
| 454-1091 | - | +++ |
| 568-1091 | +++ | - |
| Δ451-567 | +++ | - |

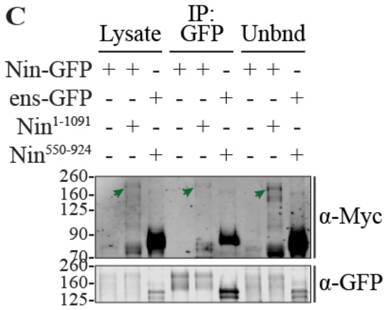


**Supplementary Figure 1.** Localization of Ninein constructs is not dependent on endogenous Ninein.
IF staining of DNA (DAPI, green) in larval fat body cells expressing Nin transgenic proteins (TagRFP fluorescence, red) in a wild-type (*Nin*$^{+}$) background.
(A) Localization patterns of TagRFP-Myc-labelled Nin transgenes using SPARC-GAL4 in a wild-type background. Multiple domains of Nin, detected with TagRFP fluorescence, localize to the MTOC. Constructs containing amino acids 454-567 localize intranuclearly.
(B) Summary of results in (A). +, weak; ++, moderate; +++, strong; -, no localization.
See Figure 2 for expression in a *Nin* null mutant (*Nin*$^{1}$) background.
Supplementary Figure 2B for fluorescence intensity profiles.
(C) Nin multimerization assay. Western blot analysis of Nin-GFP co-IP with co-overexpressed Nin$^{1\text{-}1091}$-TagRFP-Myc demonstrates that Nin can form multimers. co-IP of Nin$^{550\text{-}924}$ with ens-GFP was used as a positive control (see Figure 6B).

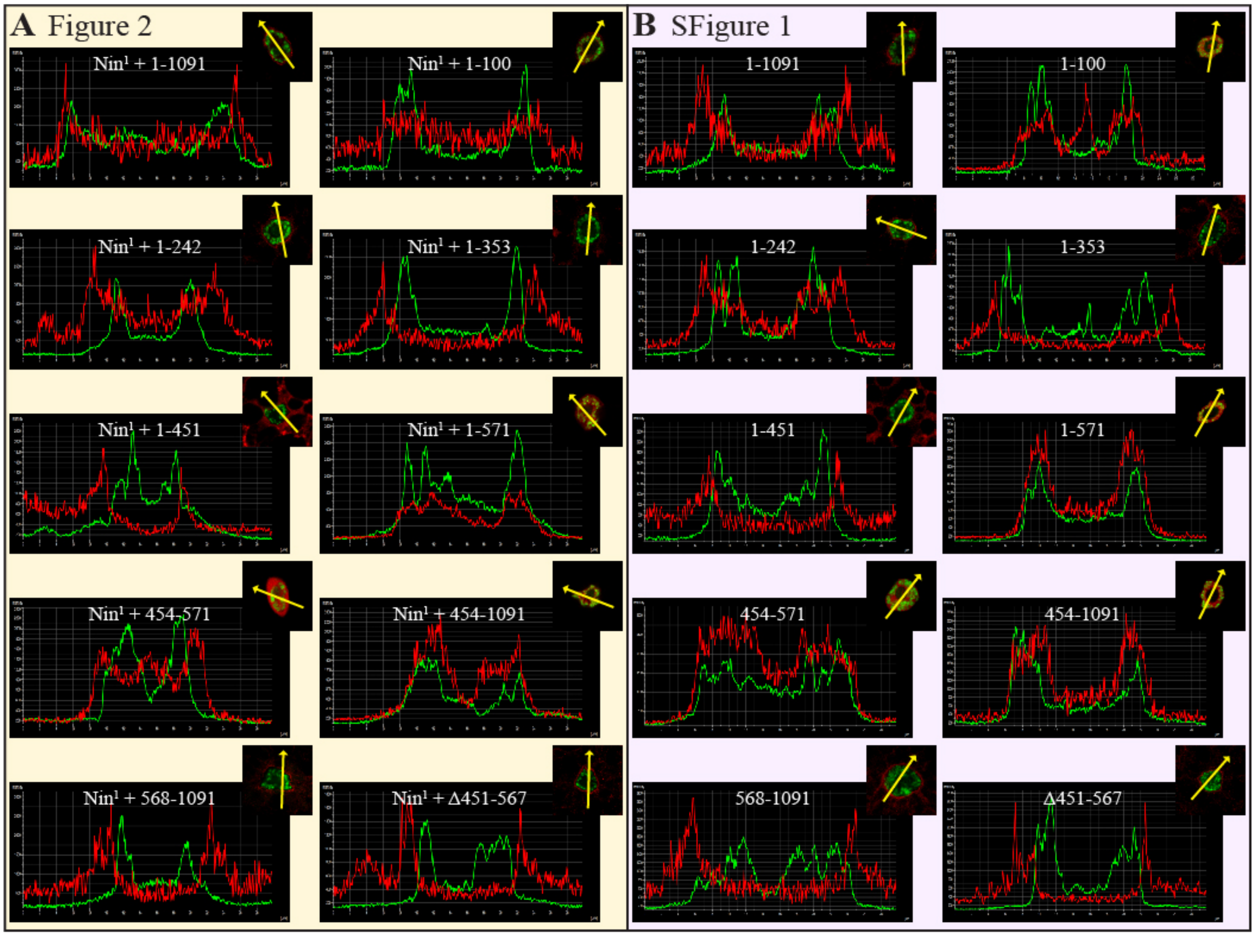


**Supplementary Figure 2.** Fluorescence intensity profiles for Figure 2 and Supplementary Figure 1.

(A-B) Inset features a single *z*-slice of images shown in Figure 2 or Supplementary Figure 1. Nuclear signal (DAPI, green), Nin signal (TagRFP fluorescence, red). Yellow arrow demarcates the *x*-axis on the intensity profile. Constructs lacking amino acids 454-567 show decreased intranuclear TagRFP levels. Amino acids 1-100 also show an enhanced intranuclear signal.

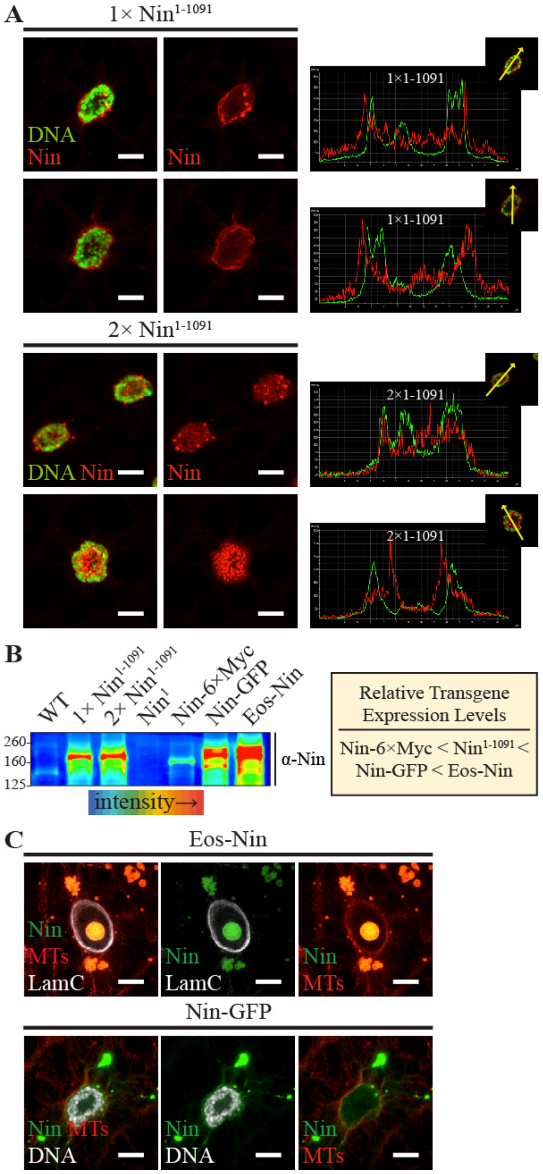


**Supplementary Figure 3.** Overexpression of Ninein increases aggregation, nuclear localization, and lethality.

(A) Localization from expression of one copy of $Nin^{1\text{-}1091}$ using SPARC-GAL4 in a wild-type background is predominantly perinuclear, weakly intranuclear, and variably produces small aggregates. Expression of two copies of $Nin^{1\text{-}1091}$ in a wild-type background results in increased aggregation and intranuclear localization. Fluorescence intensity profiles are shown on the right.

(B) Western blot analysis of fat bodies expressing Nin transgenes using an antibody against Nin. $w^{1118}$ was used as a wild-type (WT) control. Nin transgenes were driven with SPARC-GAL4. Warmer colors indicate more protein. On the right, transgenes are ranked in order of increasing relative expression levels. Overexpression of Nin-6×Myc and $Nin^{1\text{-}1091}$ are viable, whereas Nin-GFP is semi-lethal, and Eos-Nin is 100% lethal.

(C) Localization of highly expressed Eos-Nin (above) and Nin-GFP (below) transgenes using SPARC-GAL4 in a wild-type background.

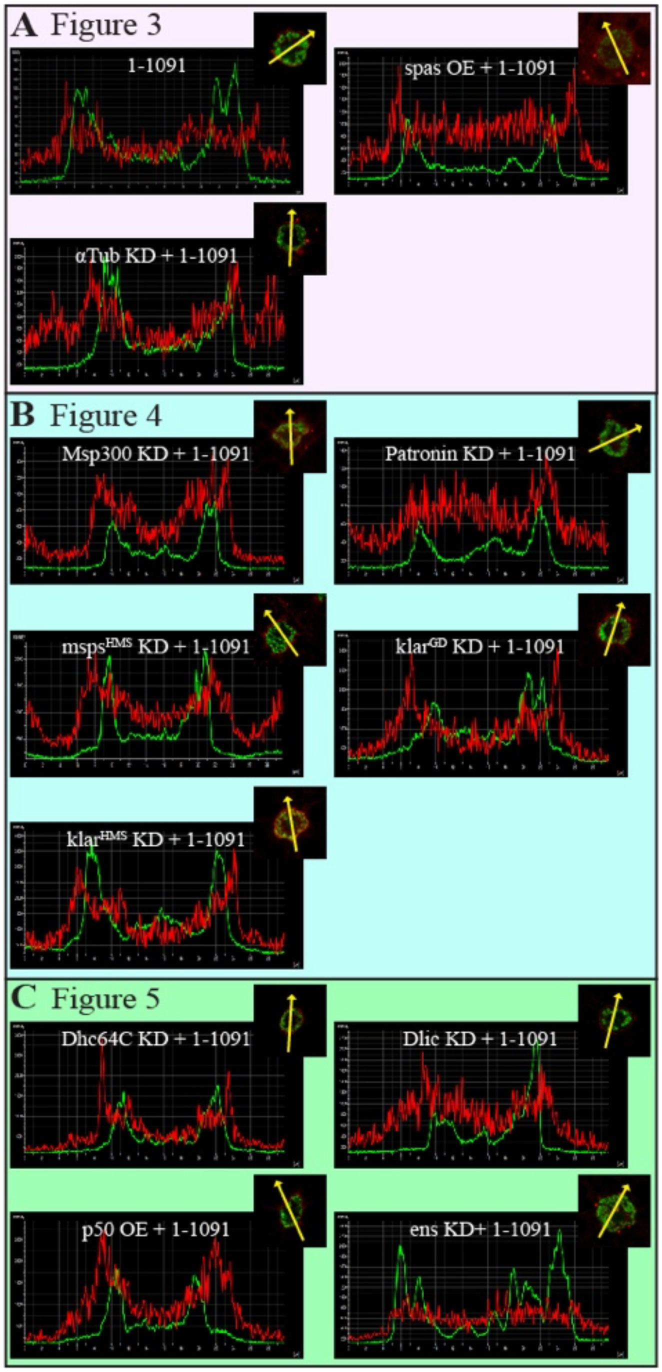


**Supplementary Figure 4.** Fluorescence intensity profiles for Figures 3-5.
(A-C) Inset features a single *z*-stack of images shown in Figures 3-5. Nuclear signal (DAPI, green), Nin signal (TagRFP fluorescence, red). Yellow arrow demarcates the *x*-axis on the intensity profile.

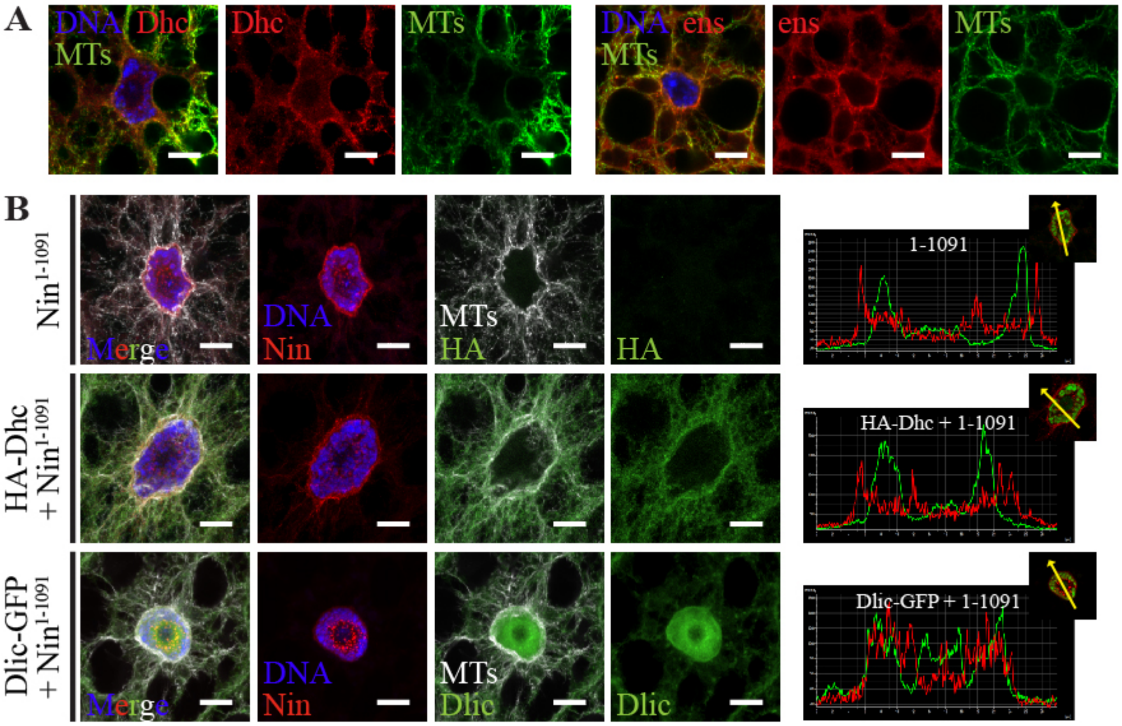


**Supplementary Figure 5.** Overexpression of Ninein and dynein does not alter microtubule organization.

(A) IF staining of wild-type larval fat body cells. DNA (DAPI, blue), Dhc (2C11-2, red) above or ens (anti-ens, red) below, and microtubules (YL1/2, green).

(B) IF staining of larval fat body cells expressing the indicated transgenes using SPARC-GAL4 in a wild-type background.

DNA (DAPI, blue), $Nin^{1\text{-}1091}$ (TagRFP fluorescence, red), Dhc (anti-HA, green) or Dlic-GFP (GFP fluorescence, green), and microtubules (YL1/2, white).

Co-overexpression of $Nin^{1\text{-}1091}$ and Dynein heavy chain (HA-Dhc) or Dynein light intermediate chain (Dlic-GFP) does not alter microtubule organization. Dlic-GFP is intranuclear and enhances Nin localization there.
